# The MarR-family regulator TilR links biosynthesis of enterotoxin tilimycin with physiologic adaptation in *Klebsiella oxytoca*

**DOI:** 10.64898/2026.11.06.765007

**Authors:** Katrin Unterhauser, André A. Grassmann, Parithi Balachandran, Anthony A. Provatas, Yanjiao Zhou, Melissa J. Caimano, Justin D. Radolf, Adam P. Matson

## Abstract

Members of the *Klebsiella oxytoca* species complex (KoSC) are ubiquitous gut colonizers that produce the DNA-damaging enterotoxin tilimycin (TM) as part of their virulence-related program. A MarR-like transcriptional regulator, herein designated TilR, controls expression of the divergently transcribed *aroX*-and NRPS-operons responsible for TM production. Deletion of *tilR* resulted in pronounced induction of both operons, elevated toxin production, and increased cytotoxicity – establishing TilR as the key repressor of TM biosynthesis. TilR is encoded downstream of the NRPS operon and binds a 20 bp motif within the *aroX*-NRPS intergenic region, spatially separated from the predicted promoters driving toxin gene expression. Analysis of global transcriptome profiles revealed that TilR, the most highly expressed of the seven MarRs in KoSC, links toxin biosynthesis to a broader transcriptional network involved in nutrient acquisition, stress adaptation, and central metabolism. In addition, TM production coincided with induction of multiple virulence-associated genes involved in adhesion, biofilm formation, and iron acquisition. Phylogenetic and structural analyses revealed that TilR is distinct from canonical MarRs and absent from members of the closely related, competing *Klebsiella pneumoniae* species complex (KpSC). These findings identify TilR as an evolutionarily specialized global regulator that coordinates TM biosynthesis with broader metabolic, stress-adaptive, and virulence-associated transcriptional responses.

**AUTHOR SUMMARY:** Members of the *Klebsiella oxytoca* species complex (KoSC) cause antibiotic-associated hemorrhagic colitis and have been linked to necrotizing enterocolitis through production of the enterotoxin tilimycin (TM). Although TM has long been recognized as a major virulence factor, little has been known about how toxin production is regulated and coordinated with broader bacterial physiology. Herein, we show that TilR acts as both a repressor of TM biosynthesis and a pleiotropic regulatory hub linking toxin production to pathways involved in nutrient acquisition, metabolism, and stress adaptation. We also found that induction of TM biosynthesis coincides with increased expression of genes associated with host colonization and survival. These observations suggest that toxin production is not an isolated event but occurs as part of a broader physiological transition that may enhance bacterial fitness in the gut. By revealing how KoSC integrates environmental signals with toxin production and other adaptive traits, this work provides a framework for understanding how common gut colonizers become disease-causing pathogens and identifies new avenues for investigating the factors that trigger toxin production *in vivo*.

## INTRODUCTION

The human gastrointestinal (GI) tract harbors diverse symbiotic bacterial species that are crucial for host physiology and immune maturation (1, 2). Preservation of gut homeostasis relies on highly efficient innate and adaptive mucosal immunity, continuous epithelial renewal, and equilibrium between resident microbial populations (3, 4). Cooperative behaviors within and between microbial species are coordinated through an array of signaling molecules and metabolites in response to nutrient flux and host-derived signals (5–7). In addition, the stable microbial community provides resistance against colonization by non-native bacteria (8, 9). However, some commensals behave as pathobionts, taking on pathogenic properties in response to changes in nutrient availability and other stress-inducing environmental perturbations (10, 11).

In recent years, members of the genus *Klebsiella* have emerged as ubiquitous gut colonizers, particularly of individuals in hospital settings (12, 13). In many cases, the environmental cues that trigger expression of virulence mechanisms and the regulators that effect this transition remain poorly understood. Members of the *Klebsiella oxytoca* species complex (KoSC) can cause pneumonia, meningitis, infections of the urinary tract and bacteremia (14). In the gut, cytotoxin-producing KoSC have been linked to antibiotic associated hemorrhagic colitis (AAHC) in younger children and adults (15, 16) and necrotizing enterocolitis (NEC) in premature infants (17–19). Both intestinal diseases are associated with treatment with β-lactam antibiotics (15, 20), which can lead to overgrowth of KoSC and production of the small molecule cytotoxins tilimycin (TM) and tilivalline (TV) (21–24). TM acts as a genotoxin by alkylating host DNA, leading to apoptosis of intestinal epithelial cells (22) and generation of pre-cancerous mutations in colonic stem cells (25). TM also exhibits antibacterial activity against multiple gut commensals (22, 24, 26) and has been linked to antibiotic resistance of luminal microbiota (24). In the presence of indole, a signaling molecule produced by gut microbiota (27, 28), TM spontaneously converts to the secondary product TV. TV stabilizes host microtubules, leading to mitotic arrest (22), and activates the host xenobiotic Pregnane X receptor (PXR), upregulating detoxifying enzymes as a means of mitigating tissue cytotoxicity (26). TM is more potent and abundant than TV, suggesting a greater role for the former in KoSC pathogenicity (22, 26, 29–31).

The biosynthesis of TM involves the conversion of aromatic substrates generated from the shikimate pathway via non-ribosomal peptide synthetases (NRPS) encoded in the *til* gene cluster (**Figure 1**). Chorismate is a branch-point metabolite essential for production of TM (23) as well as the synthesis of aromatic amino acids, vitamins, siderophores (*e.g.*, yersiniabactin and enterobactin), antibiotics, and various other aromatic products (32). The *til* gene cluster consists of the divergently oriented *aroX*-and NRPS*-*operons (21, 23, 29) separated by a 457 bp intergenic regulatory region. While the roles of most gene products encoded by these operons in TM biosynthesis are well established (23, 29), the locus also encodes a multiple antibiotic resistance regulator (MarR)-like protein that functions as a negative regulator of TM biosynthesis (33). Herein, we refer to this protein as TilR (*til* gene cluster Regulator), although it was recently designated TagR in an independent study (33). While TilR was shown to bind within the *aroX*-NRPS intergenic region (33), the mechanism by which it represses TM biosynthesis remains poorly understood, and the full extent of its regulon has yet to be defined. Of note, other transcriptional regulators (*e.g*., catabolite repression protein [CRP] (34), leucine-responsive protein [LRP] (35), integration host factor [IHF] and factor for inversion stimulation [Fis] (36) have been identified as activators of TM biosynthesis through binding to sites within the intergenic region.

**Figure 1.**
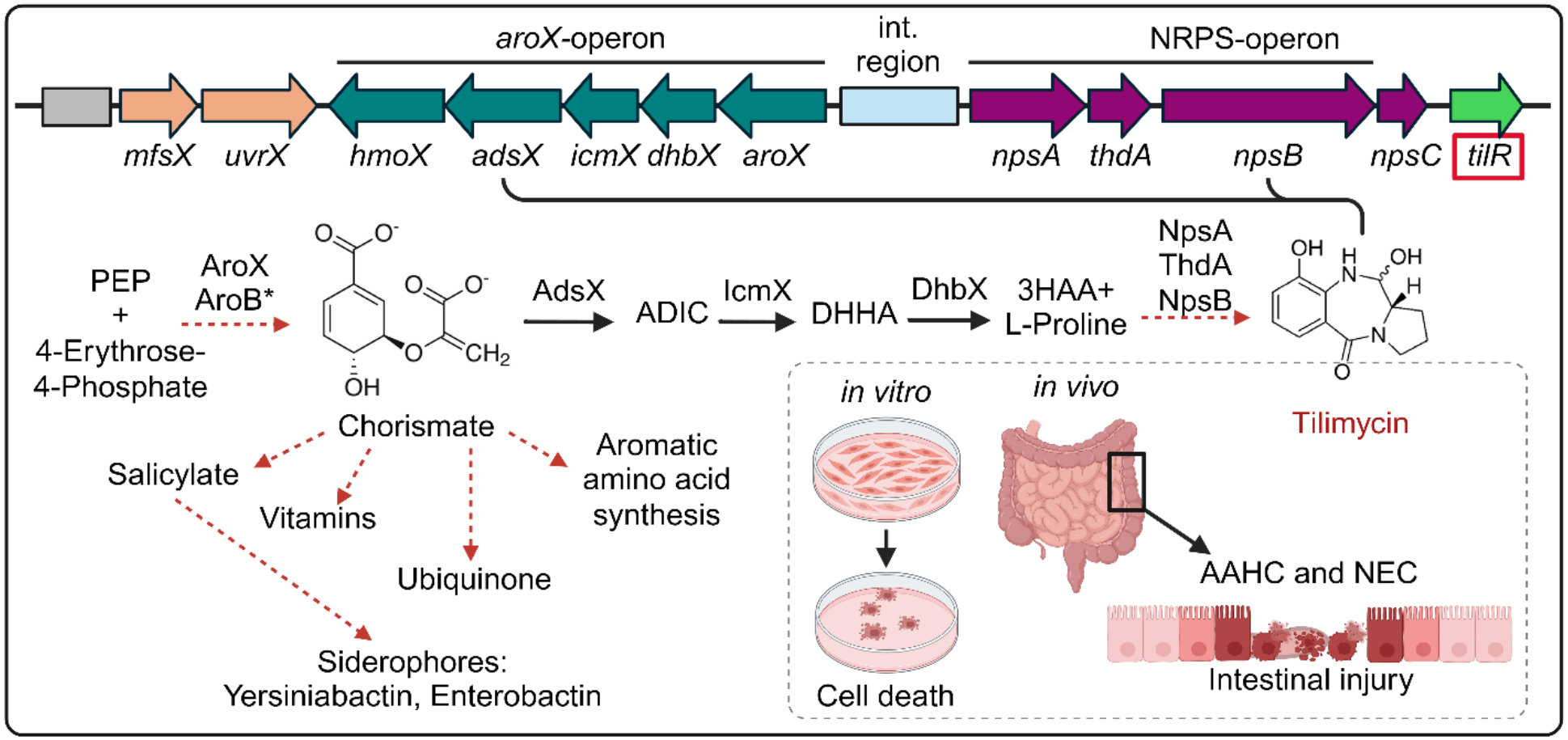
**The *til* gene cluster and the TilR-associated tilimycin (TM) biosynthetic pathway in KoSC**. The divergently oriented *aroX*-and NRPS-operons are necessary for production of TM by KoSC (23). AroX, AroB and various enzymes of the shikimate pathway (indicated by asterisk) convert phosphoenolpyruvate (PEP) and 4-erythrose-4-phosphate to the important branchpoint metabolite chorismate. Chorismate further serves as substrate for the amino-2-deoxyisochorismatase AdsX forming 2-amino-2-deoxyisochorismate (ADIC). ADIC is converted to trans-2,3-dihydro-3-hydroxyanthranilic acid (DHHA) by the isochorismatase IcmX. The oxidation of DHHA to 3-hydroxyanthranilic acid (3HAA) is catalyzed by 2,3-dihydro-2,3-dihydroxybenzoate dehydrogenase DhbX. The NpsA/ThdA/NpsB complex converts 3HAA and L-Proline (produced outside of the pathway) to the cytotoxin, TM, causally associated with antibiotic associated hemorrhagic colitis (AAHC) and necrotizing enterocolitis (NEC). Black arrows indicate where one enzyme catalyzes a reaction, whereas red arrows with dashed lines denote the involvement of multiple enzymes in a biosynthetic pathway. The *tilR* gene located downstream of the NRPS operon encodes the MarR ortholog, TilR, (red box, locus tag NDX50_RS24840, 498 bp). *npsC* encodes a polyketide antibiotic cyclase homologous to SnoaL in *Streptomyces* (133). The *mfsX* gene located downstream of the *aroX* operon encodes a major facilitator superfamily (MFS) homolog of the EmrD and MdfA multidrug efflux pumps in *E. coli* (66, 67); *uvrX* encodes a homolog of the Ecm16 DNA repair protein from *Streptomyces lasalocidi* (134). The schematic representation of the *til* gene cluster is based on the annotation of locus tags NDX50_RS24900 to NDX50_RS24840 in *K. oxytoca* AHC-6 (GenBank Accession No. NZ_CP098757.1). The grey rectangle indicates the upstream and putative regulatory region of the *mfsX-uvrX-*operon.

MarR-like regulators are a ubiquitous, ancient family of bacterial transcription factors that have evolved through extensive gene duplication, with the average bacterial genome encoding seven unique family members (37–39). MarRs from different lineages typically share <30% sequence identity and display significant structural variation, suggesting they have evolved to serve diverse regulatory functions (39). Regulation of multidrug-resistance (MDR) efflux pumps is an important and clinically relevant function of MarRs given the worldwide concern about MDR bacteria (40, 41). Control of genes involved in stress responses, catabolic pathways and virulence are additional, well-documented roles of MarR proteins (37). Genes encoding MarR proteins are frequently oriented divergently from the genes they repress or, in rare cases, activate (42). MarRs recognize palindromic DNA sequences that often overlap promoter elements, thereby blocking initiation of transcription (37). Alternatively, some MarRs bind outside of promoter regions to block transcript elongation or alter DNA topology, hindering promoter accessibility to RNA polymerase (RNAP) (42, 43). Most MarRs bind small molecule ligands such as phenolic compounds, aromatic carboxylates or antibiotics, leading to structural changes that reduce their binding affinity for DNA (44), while others dissociate from DNA following oxidation of cysteines (45). Another mechanism of gene regulation, exemplified by SlyA in *Salmonella enterica* (46) and RovA in *Yersinia enterocolitica* (47), is counter-silencing of genes repressed by histone-like nucleotide-associated protein (H-NS). By sensing chorismate-derived metabolites (**Figure 1**) produced during iron-depleted conditions, SlyA links aromatic metabolism to the control of horizontally acquired genes involved in efflux and virulence (48).

Members of the KoSC have evolved a variety of strategies to deal with competitive challenges imposed by other gut-resident pathogens including its evil cousin, *K. pneumoniae* (49–51). The production of TM presumably contributes to their success by diminishing taxonomic composition of the gut microbiota and damaging the intestinal epithelium as a means of liberating nutrients (22, 24, 26). Most bacterial toxin systems are under strict regulatory control to ensure that their production is coordinated with responses to specific environmental or nutritional signals (52). MarR-family regulators also contribute to KpSC biology. For example, KvrA regulates virulence-associated traits including capsule production (53), whereas PecS regulates urate-responsive expression of type 1 fimbriae (54). Here we show that TilR is a structurally distinct MarR-family regulator that represses TM biosynthesis and shapes global gene expression in *K. oxytoca.* While six other MarR paralogs are shared between KoSC and members of the *K. pneumoniae* species complexes (KpSC), TilR is unique to KoSC, suggesting a potential role in KoSC–KpSC competition (18, 49, 55). Our findings indicate that acquisition of the *til* gene cluster enabled KoSC to strategically link TM biosynthesis with broad physiologic reprogramming to fine-tune cellular homeostasis and enhance competitiveness within the gut lumen.

## RESULTS

### Predicted structure of TilR reveals MarR-like features

The MarR ortholog characterized herein derives from *K. oxytoca* AHC-6, a well-studied, toxigenic strain isolated from an adult with severe AAHC (21–24, 56). The *tilR* gene encodes a ∼19 kDa protein predicted by AlphaFold 3 (AF3) to consist of six α-helices and two adjacent β-strands arranged in an α1-α2-α3-α4-β1-β2-α5-α6 topology; α3, α4, β1, β2, and the connecting loop form the characteristic MarR winged helix-turn-helix (wHTH) motif (42, 57) (**Figure 2A**). Like other characterized MarRs (42), TilR is predicted to form a triangular homodimer through interactions between residues in α1, α5 and α6 (**Figure 2B**). When TilR is modeled in complex with the *aroX*-NRPS intergenic region (**Figure 1**), AF3 predicts that, as shown for other canonical MarRs (58), α4 and α4’ insert into adjacent DNA major grooves while the loops of the wHTH motifs fit within the neighboring minor grooves (**Figure 2B**). Analysis of electrostatic potential shows a strong, positively charged surface for DNA binding (**Figure 2C**). Residues K79 and R82 in α4 and R100 in the wHTH motif are predicted to contribute to DNA binding by their respective positioning in the major and minor grooves (**Figure 2D**).

**Figure 2.**
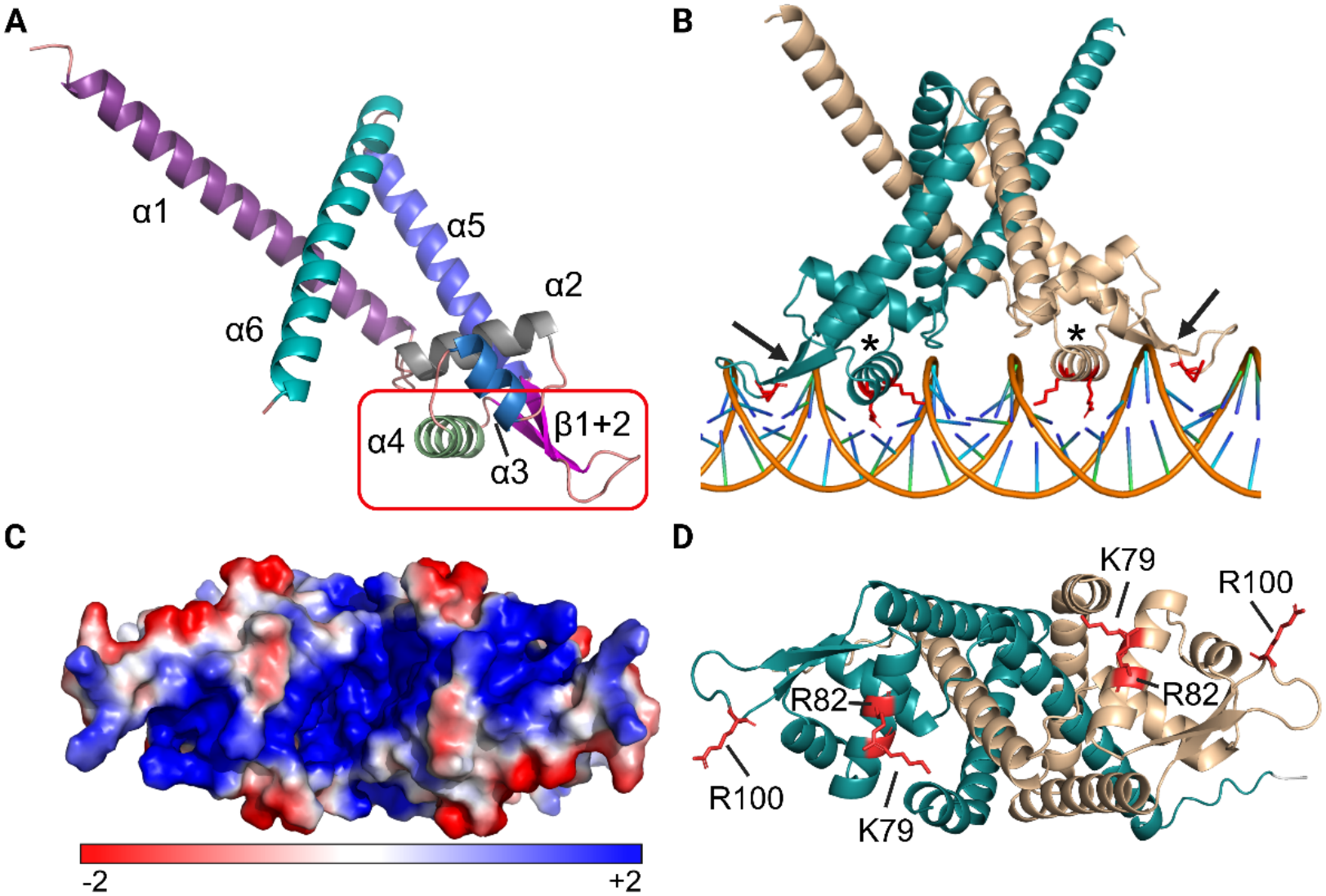
Structural models of the TilR monomer and DNA bound dimer. (**A**) AF3 predicts that the *K. oxytoca* TilR monomer consists of six α-helices (α1-α6) and two β-sheets (β1 and β2) with α3, α4, β1, β2, and the connecting loop forming a characteristic winged helix-turn-helix (wHTH) motif crucial for DNA binding (red box). (**B**) AF3 model of the dimer (monomers in teal and beige) complexed with double-stranded DNA from the *aroX*-NRPS intergenic region. The α4 and α4′ helices (indicated by asterisks) are precisely positioned within two adjacent major grooves while the loop of the wHTH motif fits within the neighboring minor grooves (arrows). (**C**) Electrostatic surface potential of the homodimer (visualized using PyMOL and rotated 90° from its orientation in B) shows the positively (blue) charged surface predicted to reside at the interface for DNA binding. (**D**) Putative DNA-binding residues K79, R82 and R100 (depicted in red in each monomer) fitting into the DNA grooves.

### TilR represses production of TM

To investigate whether TilR acts as a regulator of TM production, we generated a scarless chromosomal *tilR* deletion in *K. oxytoca* AHC-6 (Δ*tilR*; see Materials and Methods and **Figure S1**) and verified that loss of TilR had no effect on growth in LB + 2.5 g/L glucose (LBG) (**Figure 3A**). By mass spectrometry, TM was detected initially at 6 h in WT culture supernatants and further increased at 8 h, showing that toxin production is growth dependent as previously described (23). Δ*tilR*, in contrast, began producing TM as early as 4 h and produced markedly greater amounts of TM compared to WT at 6 and 8 h (**Figure 3B**). A cytotoxicity assay confirmed that filtered supernatants from Δ*tilR* caused significantly greater toxicity towards HT-29 human colon cancer cells compared to WT (**Figure 3C**). Complementation of Δ*tilR* was achieved with p*tilR*, the low copy plasmid pAR181 containing *tilR* downstream of the *aroX*-NRPS intergenic region (**Figures 1 and S2A**) on the assumption that the intergenic region contains the promoter driving expression of *tilR* as well as the NRPS operon. Compared to WT, the complemented strain displayed similar growth (**Figure S2B**) and generated comparable concentrations of TM in culture supernatants (**Figure S2C**). Together these findings confirmed that TilR is a repressive regulator for TM biosynthesis.

**Figure 3.**
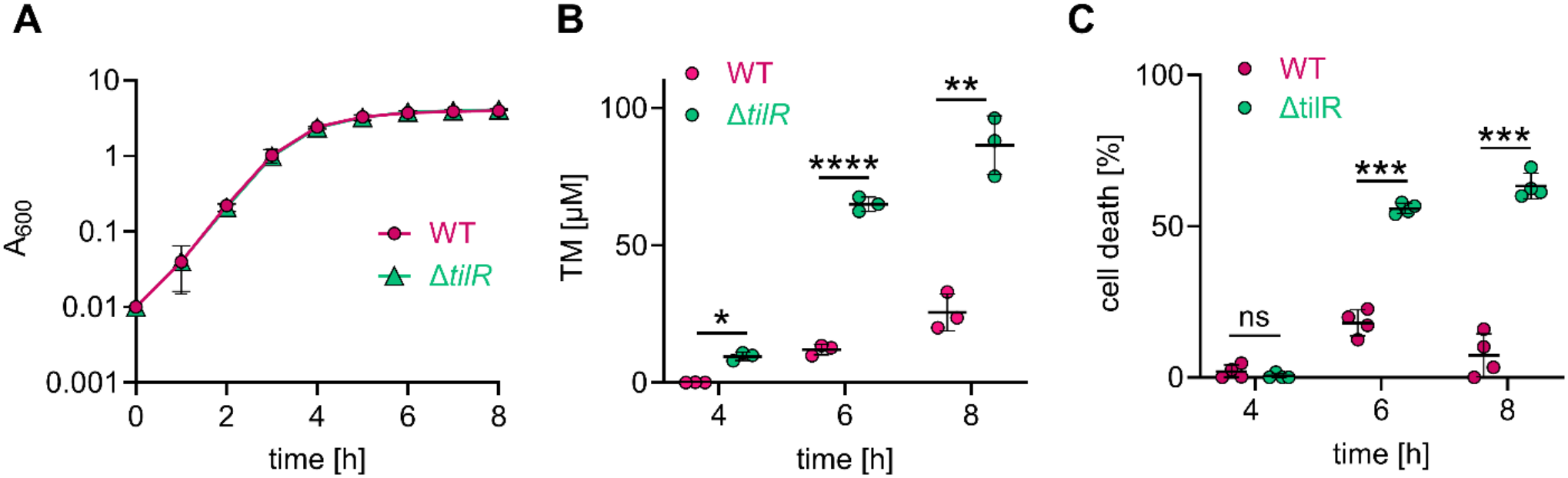
TilR represses production of TM. (**A**) Growth curves of *K. oxytoca* AHC-6 WT and Δ*tilR* in LB + 2.5 g/L glucose (LBG) (mean ± SD, n=3). (**B**) TM concentrations in culture supernatants (mean ± SD, n=3) measured by MS at 4, 6 and 8 h of growth in LBG. (**C**) Toxicity of 4, 6 and 8 h culture supernatants (1/2 dilution) for HT-29 colon cancer cells (mean ± SD, n=3). Statistical analysis for B and C was performed using Two-way repeated measures (RM) Anova followed by Sidaks multiple comparison (*P≤0.05, **P≤0.01, ***P≤0.001, **** P≤0.0001).

### TilR binds to the *aroX*-NRPS intergenic region

Genetic loci regulated by MarRs typically consist of divergently oriented genes/operons encoding the MarR homolog and its target gene(s) separated by an intergenic region (42, 59). This configuration allows the transcription factor to bind to the intergenic region, regulating both its own expression and the activity of the divergently oriented gene(s). Despite its atypical location downstream of the *til* gene cluster (**Figure 1**), TilR’s role as a repressor of toxin production led us to examine whether TilR binds within the *aroX*-NRPS intergenic region. As an initial approach, we overexpressed and purified recombinant His-tagged TilR from *Escherichia coli* for use in Electrophoretic Mobility Shift Assays (EMSAs). However, during size-exclusion chromatography, recombinant His-tagged TilR consistently eluted as a tetramer deemed unlikely to be functional (60). As an alternative, we performed DNA pull-down assays (described below) (61) and, to this end, created p*tilR*-HA encoding TilR containing a C-terminal hemagglutinin (HA) tag. Repression of TM production during early growth of *K. oxytoca* AHC-6 Δ*tilR* harboring p*tilR*-HA confirmed that TilR-HA is functional (**Figure S3A**). Notably, TilR was barely detectable at early time points (**Figure S3B**) when it acts as a repressor of TM biosynthesis in its presumptive unliganded state.

To enhance TilR levels for DNA pull-down assays at early time points, we moved TilR-HA to a plasmid under the control of an IPTG-inducible promoter, generating pIPTG-*tilR*-HA **(Figure S3C)**. Induction of this construct led to substantially greater expression of TilR-HA during early growth (**Figure S3D**). Biotinylated DNA fragments spanning most of the *aroX*-NRPS intergenic region (448 bp), as well as truncated fragments (**Figure 4A**), were coupled to streptavidin beads and used to pull-down TilR-HA from whole-cell lysates prepared from the *trans*-complemented strain 2 h after IPTG induction (**Figure 4B**). The 448 bp *aroX*-NRPS intergenic fragment (fragment #1) successfully captured TilR-HA, whereas no bands were detected using a 255 bp truncated fragment upstream of *aroX* (fragment #2) or a 191 bp truncated fragment upstream of *npsA* (fragment #3) (**Figures 4A and C**). A 100 bp sequence located in the middle of the intergenic region (fragment #4) also efficiently captured TilR-HA (**Figures 4A and C**), narrowing the TilR binding site to this region. MarR family proteins typically bind to imperfect inverted palindromic sequences of approximately 20 bp (42). To identify potential TilR binding motifs, the 100 bp fragment was analyzed using the Emboss palindrome finder (62). Within this fragment, we identified a 20 bp sequence containing a 6 bp perfect inverted repeat (underlined), TGATTTTTTTTTGGAAATCA (**Figure 4A**). Notably, this sequence corresponds to the binding site recently identified by Perez *et al.* (33). A mutated version of this inverted repeat (fragment #4a; GTCGCGTTTTTTGGCGCGAC) failed to capture TilR-HA (**Figure 4C**), confirming this sequence as the TilR binding site.

**Figure 4.**
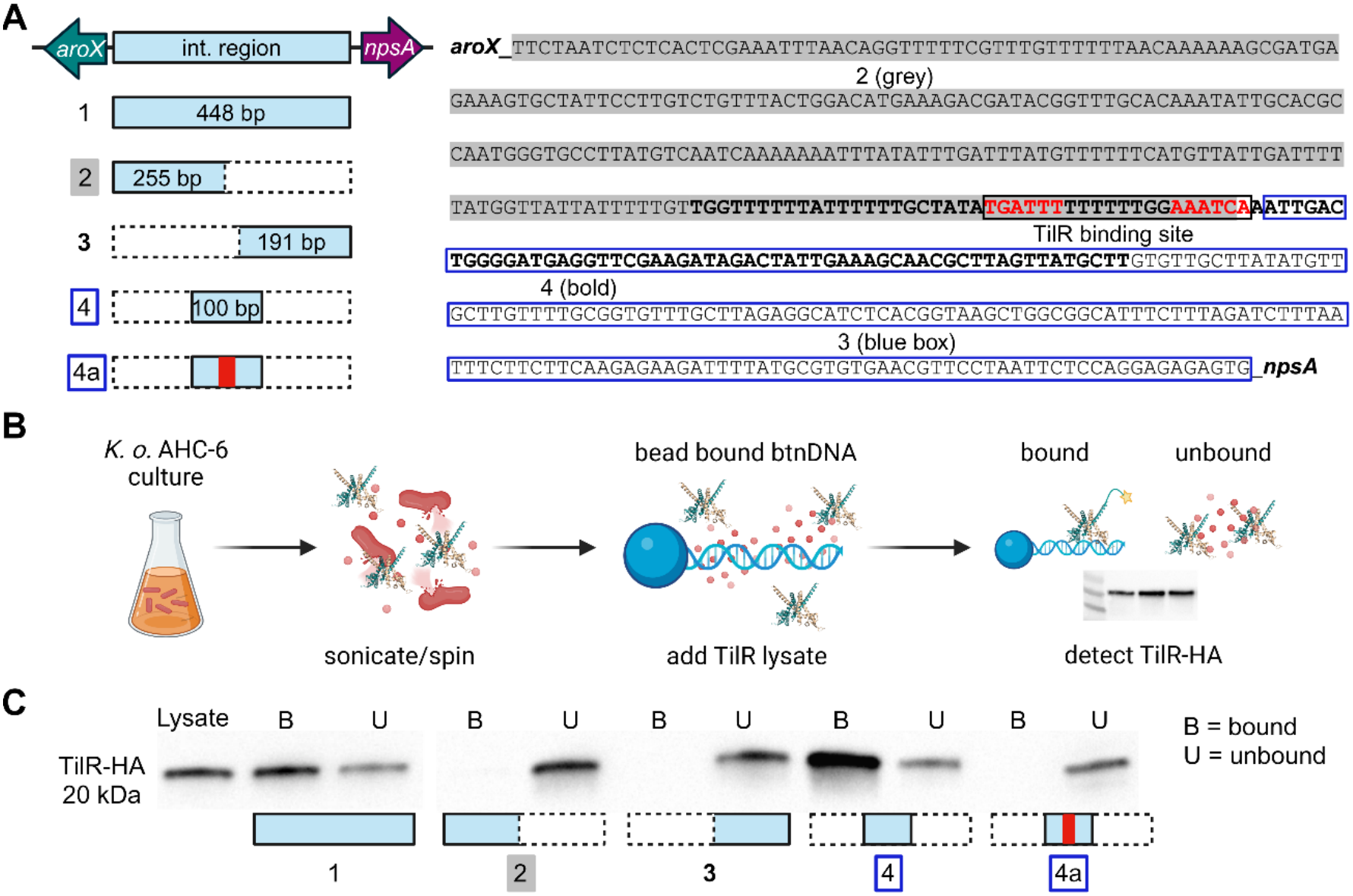
Identification of the TilR binding site in the *aroX-*NRPS intergenic region. (**A**) Schematic of biotinylated fragments used for DNA pull-down assays. Fragment 1 (448 bp) spans most of the entire intergenic region; fragments 2 (255 bp) and 3 (191 bp) cover regions upstream of *aroX* and *npsA*, respectively. Fragment 4 (100 bp) lies centrally, while 4a is the same fragment containing a mutated TilR binding site (see below). The sequence on the right displays the aroX-NRPS intergenic region (5’-3’) with the fragments distinctively labeled and the 6bp repeats of the TilR binding site displayed in red. (**B**) Workflow of the pull-down assay used to identify the TilR binding site. *K. oxytoca* AHC-6 Δ*tilR* harboring pIPTG-*tilR*-HA was induced for 2 h with 100 µm IPTG. Cells were pelleted, sonicated and spun again to generate whole cell lysates. Cell lysates were incubated with biotinylated fragments 1, 2, 3, 4, and 4a coupled to Streptavidin magnetic beads. The supernatant was separated from beads, and the beads were further washed with PBS to remove any unbound TilR prior to SDS-PAGE. Immunoblots were performed to detect TilR-HA in input lysate, bound to beads (B, fractions pulled down with beads) and in supernatants (unbound, U). (**C**) Representative immunoblot images (n=3) of bound and unbound TilR-HA along with schematics of the biotinylated DNAs used for the pull-downs. Fragment 4 contains the predicted TilR binding motif (TGATTTTTTTTTGGAAATCA) while 4a carries the mutated motif (GTCGCGTTTTTTGGCGCGAC) with the 6 bp that were altered highlighted in red.

### TilR represses bidirectional transcription of toxin genes without directly occluding predicted promoters

Given the location of the TilR binding site within the *aroX*-NRPS intergenic region, we next sought to better understand how TilR coordinately represses transcription of both operons. Accordingly, we performed RNA-seq on WT and Δ*tilR* strains to examine transcriptional activity across the *aroX*-NRPS intergenic regulatory region at 3 and 6 h of growth in LBG, corresponding to periods before and during TM synthesis in WT, respectively (**Figures 3A and B**). This analysis demonstrated that all genes of the *aroX* and NRPS operons were repressed by TilR at 3 h but not at 6 h (**Figure 5A and B, Table S3**), confirming that TilR bidirectionally represses transcription. We next analyzed strand-specific directional read coverage, which revealed a 202 bp interval lacking detectable RNA-seq coverage within the 457 bp *aroX*-NRPS intergenic region. This region was flanked by sharply defined, oppositely oriented reads consistent with divergent transcription start sites (TSSs) for the *aroX*-and NRPS-operons (**Figure 6A**). Guided by these data, a promoter prediction algorithm (63) identified candidate promoters for both *aroX* and *npsA* outside the TilR binding site (**Figure 6B**), indicating that TilR is unlikely to repress transcription through direct occlusion of RNAP-promoter interactions. Binding sites for four activators – CRP (34), LRP (35), Fis, and IHF (36) – have been mapped to this region; of these, only the LRP site upstream of *npsA* overlaps the TilR motif (**Figure S4**).

**Figure 5.**
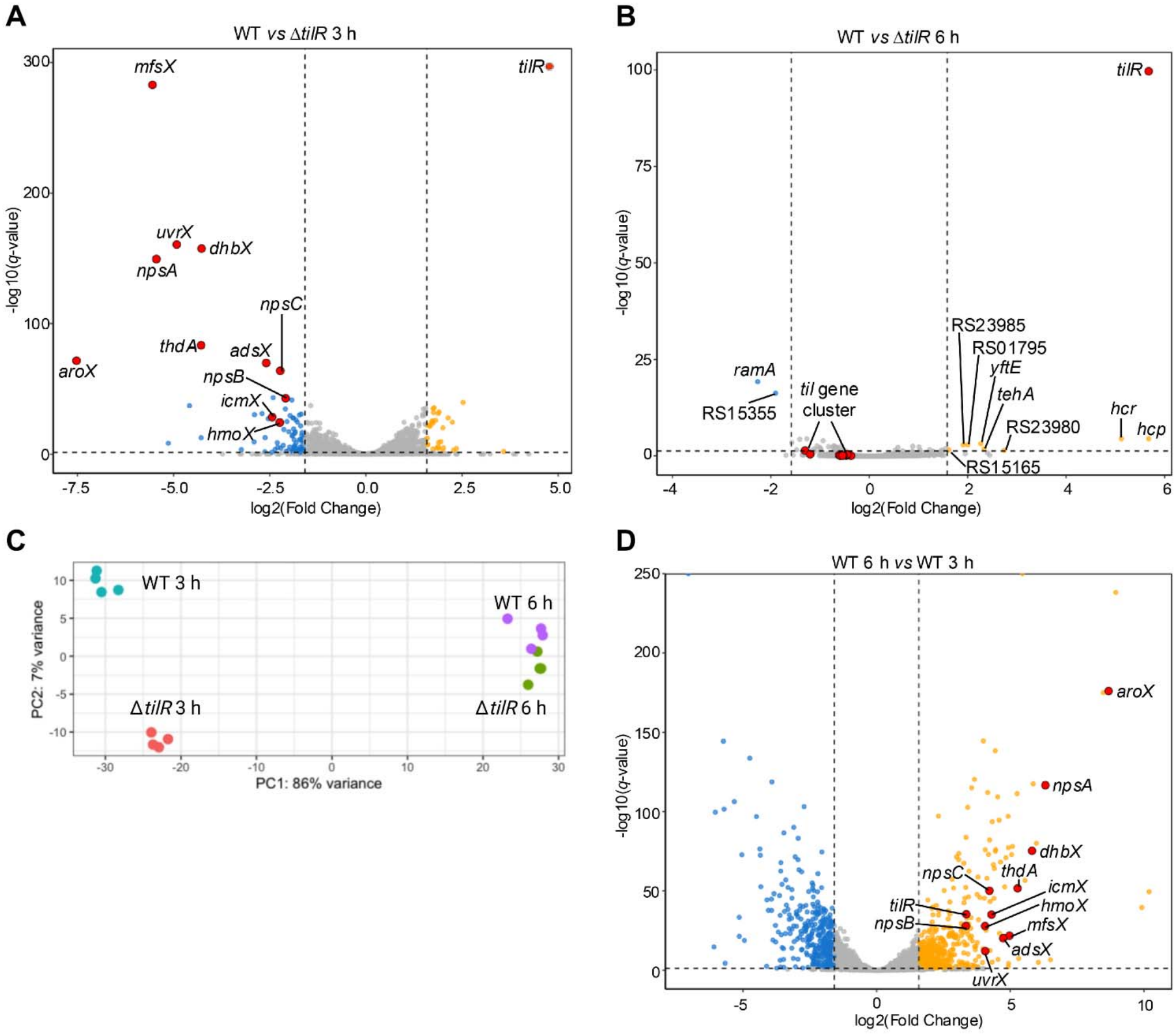
TilR functions as a global regulator predominantly during early growth. Volcano plots depicting fold changes in transcript levels for WT vs Δ*tilR* at 3 h (**A**) and 6 h (**B**). **(C)** Principal component analysis (PCA) plot comparing the transcriptomes of *K. oxytoca* AHC-6 WT and Δ*tilR* at 3 h and 6 h of growth in LBG (based on analysis of four biological replicates). (**D**) Volcano plot depicting fold changes in transcript levels for WT at 6 h vs WT at 3 h. Genes not differentially expressed are shown in grey. Repressed and activated genes are shown in blue and yellow, respectively. Members of the *til* gene cluster are highlighted in red. Additional notable genes discussed in the main text are labeled by gene name or abbreviated locus tags (from NDX50_RSXXXX to RSXXXX).

**Figure 6.**
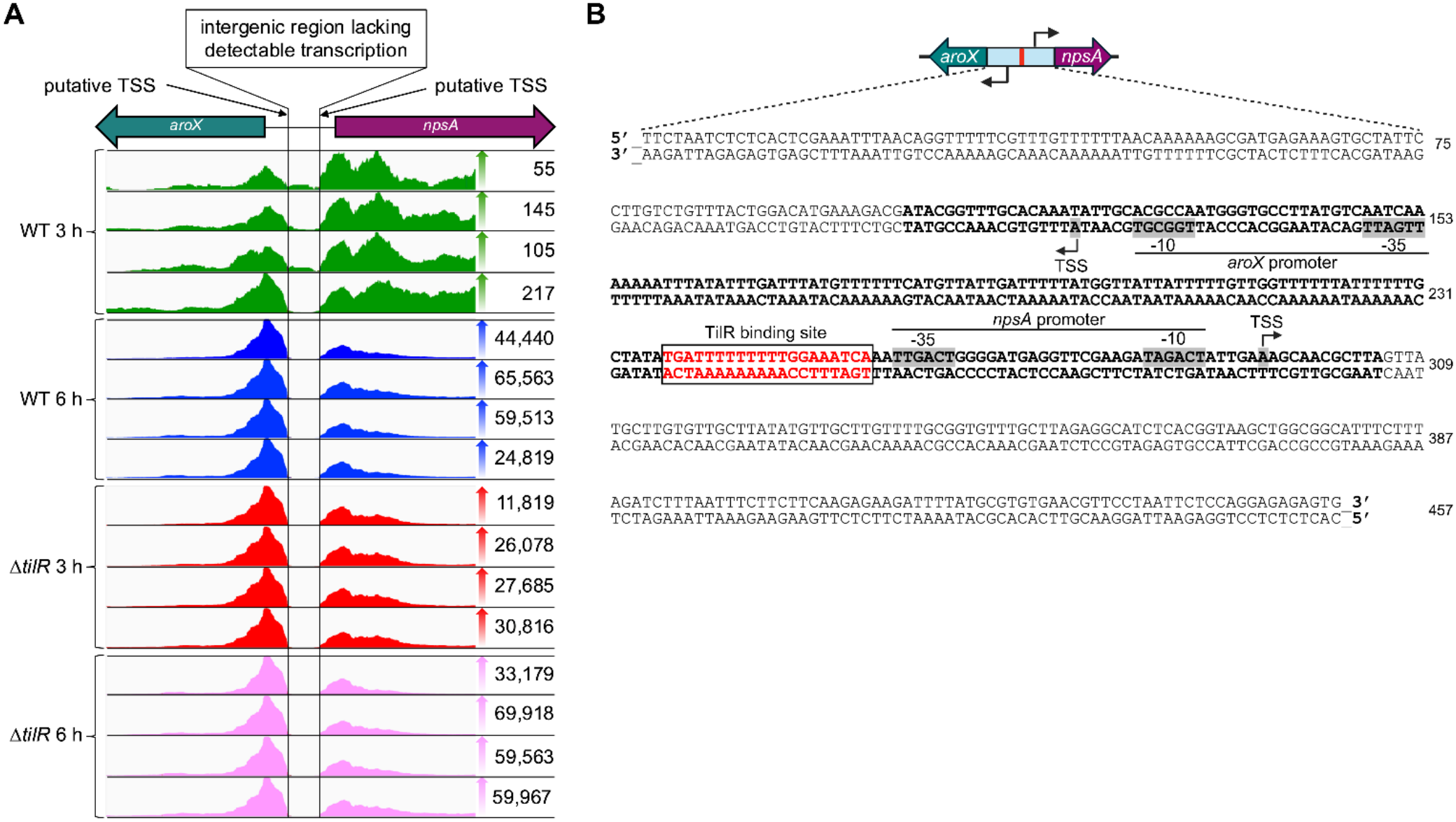
RNA-seq analysis of the *aroX*-NRPS intergenic region enables identification of candidate promoters driving bidirectional transcription of TM biosynthesis genes. (A) Read coverage across the *aroX*-NRPS intergenic region is displayed with reads aligned in the direction of *aroX* (left) and to *npsA* (right). Each horizontal track represents an individual RNA-seq library, while colored histograms indicate relative read depth; the number to the right of each histogram indicates the maximum per-base read depth within the displayed interval. The vertical black line marks the portion of the intergenic region devoid of mapped reads across all conditions examined in WT and Δ*tilR*. (B) Schematic representation of the entire *aroX*–NRPS intergenic region (in 5′–3′ and 3′–5′ orientation) with nucleotide numbers indicated to the right of the sequence. The transcriptionally silent fragment is displayed in bold letters, while the TilR binding site is highlighted in red. Promoters for *aroX* and *npsA* were predicted using the web based Neural Network Promoter prediction tool (63) and are highlighted in grey (-10 and-35 region); transcriptional starts sites (TSS) are indicated with an arrow in the direction of transcription. The localization of both predicted TSSs within the region lacking detectable RNA-seq coverage may reflect incomplete representation of transcript 5’ ends due to processing or technical limitations of conventional RNA-seq.

### TilR functions as a global regulator in addition to repressing the *til* gene cluster

MarR-like transcriptional factors traditionally have been thought of as sensors of toxic metabolites and regulators of efflux pumps for small molecules and antibiotics (40, 64, 65). Many, however, have been shown to regulate a broad range of metabolic pathways and signaling networks, including those involved in catabolism of aromatic compounds and responses to stress (37, 42). We, therefore, used RNA-seq to assess TilR’s genome-wide transcriptional impact in *K. oxytoca* AHC-6. WT and Δ*tilR* transcriptomes were analyzed at 3 and 6 h of growth in LBG, as described above. Principal component analysis (PCA) revealed clear separation between WT and Δ*tilR* transcriptomes at 3 but not 6 h (**Figure 5C**), indicating that TilR exerts broad regulatory activity primarily during early growth. Accordingly, 120 genes were differentially expressed (91 repressed, 29 activated) at 3 h compared to only 10 (8 activated, 2 repressed) at 6 h. Consistent with the operon-specific analyses described above, genes within the *til* gene cluster, including the *aroX,* NRPS, and neighboring *mfsX-uvrX* operons (**Figure 1**), were repressed by TilR at 3 h but not at 6 h (**Figure 5A and B; Table S3**). Notably, *mfsX,* which encodes a predicted major facilitator superfamily transporter (66, 67), was repressed 47-fold by TilR, comparable to the magnitude of repression observed for *npsA* (44-fold) and second only to *aroX* (184-fold) (**Figure 5A**). Outside the *til* gene cluster, the most pronounced change was activation of the *hcp-hcr* operon (∼43-fold at 6 h) (**Figure 5B**), involved in oxidative and/or nitrosative stress responses (68, 69). Most other TilR-regulated genes displayed more modest (3-5-fold) changes (**Table S4**). Because certain MarR-family members regulate gene expression by counteracting H-NS-mediated repression (46, 47), we considered whether TilR functions through a similar counter-silencing mechanism. However, we observed no TilR-dependent changes in the expression of *wzi* (NDX50_RS25340) and *wabG* (NDX50_RS06795), two H-NS-regulated genes in *K. pneumoniae* (70).

#### Genes regulated by TilR at 3 h

The 91 genes repressed by TilR encode proteins involved in diverse physiologic processes (**Figure S5A; Table S4)**; functional themes included control of efflux pumps and transporters, nutrient assimilation and response to stress, along with altered expression of regulatory proteins involved in central metabolism and oxidative stress. The 29 activated genes primarily encode proteins involved in protein synthesis (translation), transport of sugars and general cellular processes (**Figure S5A; Table S4)**. All gene locus tags indicated below are abbreviated from the full format NDX50_RSXXXX to RSXXXX for simplicity.

MarRs are well-known regulators of efflux pumps and transport proteins (39). In addition to *mfsX,* TilR-repressed genes included those encoding a second multidrug efflux facilitator superfamily (MFS) transporter (RS23885) homologous to the *E. coli* multidrug efflux transporters EmrD and MdfA (66, 67) (**Figure S6A**, **Table S4**). Other repressed efflux-associated genes encode an AcrZ family multidrug efflux pump-associated protein (RS16585) that modulates the AcrAB-TolC system involved in antibiotic resistance in *E. coli* (71, 72), as well as a TolC family protein (RS01880) homologous to *E. coli* CusC, an outer membrane component of a heavy metal efflux pump (73). TilR additionally repressed eight genes encoding proteins predicted to function in sugar transport, including a substrate-binding protein (*rbsB*, RS07725) and an ATP-binding subunit (RS15795), both of which are components of ATP-binding cassette (ABC) transporters; a phosphotransferase system (PTS) fructose transporter subunit IIC (RS04545); two glycoside-pentoside-hexuronide (GPH) cation symporters of the MFS family (RS20545 and RS04360); and a tripartite ATP-independent periplasmic transporter (TRAP) subunit (RS13060) (**Figure S6A**, **Table S4**). Additional TilR-repressed genes encoded transporters for diverse substrates, including a MFS-type 3-(3-hydroxy-phenyl) propionate transporter (*mhpT,* RS23180), the glycine betaine/L-proline ABC transporter ATP-binding protein ProV (*proV,* RS01425) (74), xanthine/uracil transporters (RS14535 and RS17875), and the permease subunit of the phosphonate ABC transporter (*phnE,* RS13325). Three repressed genes encode putative leader peptides involved in the regulation of amino acid biosynthesis (*ilvL* [RS08375]*, hisL* [RS25160]*, thrL* [RS11485]). In addition, TilR repressed the expression of genes associated with fatty acid and polyketide biosynthesis (*acpP,* RS18435) (75), adaptation to nutrient starvation (*rmf,* RS17620) (76), and oxidative/nitrosative stress responses (*yftE,* RS10080) (77).

Five genes repressed by TilR encode transcriptional factors known to regulate diverse physiologic processes in other bacteria; this finding suggests that TilR also regulates genes indirectly by integrating within other transcriptional networks. TilR repressed a GntR family transcriptional regulator (*pdhR,* RS12180) of the pyruvate dehydrogenase (PDH) multienzyme complex. In *E. coli*, PdhR plays a key role sensing pyruvate formed during glycolysis and repressing genes needed for oxidative metabolism when pyruvate levels are low (78, 79). TilR also repressed a MerR-family regulator (RS17995) that shares homology with the *E. coli* regulators SoxR, CueR and ZntR, which respond to oxidative stress and heavy metals (80, 81), and a TraR-family protein (RS16875), which is predicted to participate in maintaining membrane integrity under periplasmic stress conditions in *E. coli*. (82, 83). The repressed AraC-(RS16385) and LysR-(RS18380) family regulators (**Figure S6A**, **Table S4**) belong to two of the most ubiquitous families of bacterial transcriptional regulators (84, 85). These regulators can function as activators, repressors, or both, and typically regulate genes involved in carbon or nitrogen metabolism, stress response, motility or virulence.

#### Genes regulated by TilR at 6 h

As shown in the PCA plot (**Figure 5C**), the transcriptomes of WT and Δ*tilR* converge at 6 h suggesting that TilR’s regulatory function is for the most part absent during stationary phase. Nevertheless, 10 genes were differentially expressed at this late time point (**Figures 5B and S5B**). Five of the activated genes encode proteins that protect against oxidative and/or nitrosative stress. The bicistronic operon *hcp* (RS17255) and *hcr* (RS17250) demonstrated the highest folds of activation (**Figure 5B, Table S4**). In *E. coli*, the hybrid cluster protein, Hcp, functions as a high affinity nitric oxide (NO) reductase during anaerobic growth (68), while Hcr is a NADH-dependent Hcp reductase (68). Hcp also exhibits oxygen-sensitive hydroxylamine reductase activity (86). The aforementioned *yftE* is predicted to repair iron-sulfur-containing proteins exposed to oxidative/nitrosative stress (77), whereas RS01795 encodes a YgbA family protein predicted to play a role in defense against reactive nitrogen species (87). *tehA* (RS20275) encodes a tellurite resistance protein upregulated in response to NO in *E. coli* (88). The two genes repressed at the late timepoint comprise a bicistronic operon encoding the regulator *ramA* (RS15360) and a metallo-β-lactamase (RS15355) (**Figure 5B, Table S4**). In *K. pneumoniae,* expression of RamA is induced by a variety of ligands, including chemotherapeutic phenothiazines (89), and modulates virulence by activating genes involved in antibiotic resistance (90) and lipid A biosynthesis (91).

#### Identification of a predicted TilR binding motif

We next used the *aroX*-NRPS intergenic TilR binding motif (**Figure 4A**) to search for additional binding sites in *K. oxytoca* AHC-6. Upstream regions of the highly repressed *mfsX*-*uvrX* operon were initially examined, and a putative TilR binding motif was identified upstream of *mfsX* (TGATTTCTTTTTTGCAATTA). To identify additional potential TilR targets, binding motifs derived from these two sites were used to scan upstream regions of the 130 genes that were differentially expressed in Δ*tilR* at 3 and 6 h. This analysis identified TilR binding sites upstream of 20 genes encoding proteins with diverse predicted functions, as well as several hypothetical proteins, whose fold changes in transcript levels were substantially lower than those of *aroX*, *npsA,* and *mfsX* (**Table S5**). Based on the 22 identified binding sequences, a consensus TilR binding motif was generated: TGATTTWKYTTTBGMAATYA (**Figure S7**).

#### TM production in WT K. oxytoca coincides with induction of TilR-independent virulence-associated genes

Our RNA-Seq dataset enabled assessment of transcriptional changes in WT *K. oxytoca* coinciding with the transition from non-toxin-producing (3 h) to toxin-producing (6 h) growth conditions (**Figure S5C; Table S6**). As expected, genes of the *til* cluster including the *mfsX-uvrX* operon were strongly induced at 6 h, when TilR is presumably ligated and dissociated from its binding site (**Table 1**, **Figure 5D**). Using a recent dataset of known virulence factors in *K. pneumoniae* (92), we identified 16 TilR-independent, virulence-associated homologs that were induced concomitantly with TM biosynthesis genes (**Table 1, Figure S6B**). Notably, these included genes involved in adhesion and biofilm formation, allantoin utilization, and iron acquisition (yersiniabactin synthesis).

**Table 1.**
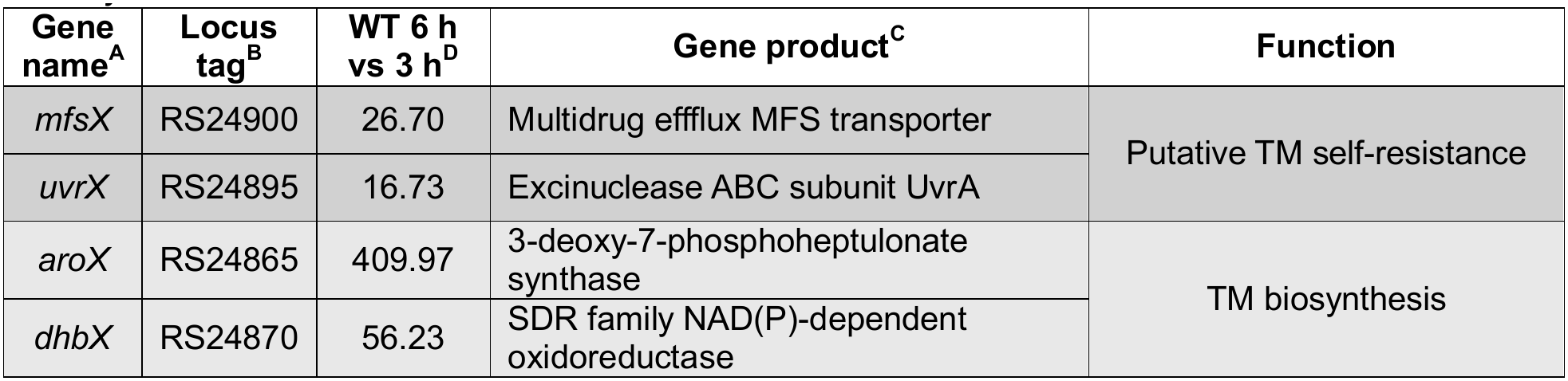
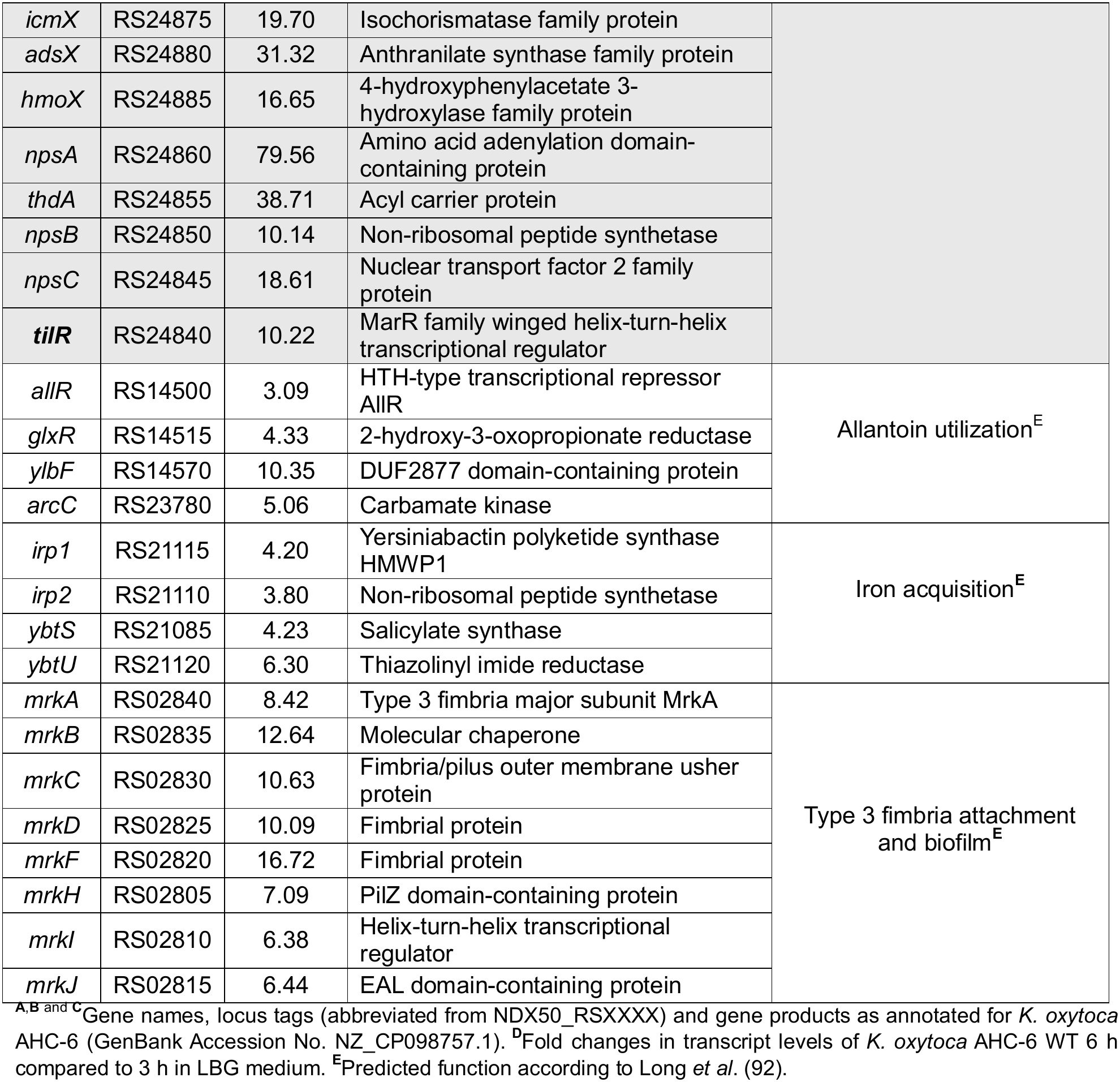
Induction of the *til* gene cluster and TilR-independent virulence-associated genes in WT *K. oxytoca* AHC-6.

| **Gene name<sup>A</sup>** | **Locus tag<sup>B</sup>** | **WT 6 h vs 3 h<sup>D</sup>** | **Gene product<sup>C</sup>** | **Function** |
|---|---|---|---|---|
| *mfsX* | RS24900 | 26.70 | Multidrug effflux MFS transporter | Putative TM self-resistance |
| *uvrX* | RS24895 | 16.73 | Excinuclease ABC subunit UvrA |  |
| *aroX* | RS24865 | 409.97 | 3-deoxy-7-phosphoheptulonate synthase | TM biosynthesis |
| *dhbX* | RS24870 | 56.23 | SDR family NAD(P)-dependent oxidoreductase |  |
| *icmX* | RS24875 | 19.70 | Isochorismatase family protein |  |
| *adsX* | RS24880 | 31.32 | Anthranilate synthase family protein |  |
| *hmoX* | RS24885 | 16.65 | 4-hydroxyphenylacetate 3-hydroxylase family protein |  |
| *npsA* | RS24860 | 79.56 | Amino acid adenylation domain-containing protein |  |
| *thdA* | RS24855 | 38.71 | Acyl carrier protein |  |
| *npsB* | RS24850 | 10.14 | Non-ribosomal peptide synthetase |  |
| *npsC* | RS24845 | 18.61 | Nuclear transport factor 2 family protein |  |
| ***tilR*** | RS24840 | 10.22 | MarR family winged helix-turn-helix transcriptional regulator |  |
| *allR* | RS14500 | 3.09 | HTH-type transcriptional repressor AllR | Allantoin utilization<sup>E</sup> |
| *glxR* | RS14515 | 4.33 | 2-hydroxy-3-oxopropionate reductase |  |
| *ylbF* | RS14570 | 10.35 | DUF2877 domain-containing protein |  |
| *arcC* | RS23780 | 5.06 | Carbamate kinase |  |
| *irp1* | RS21115 | 4.20 | Yersiniabactin polyketide synthase HMWP1 | Iron acquisition<sup>**E**</sup> |
| *irp2* | RS21110 | 3.80 | Non-ribosomal peptide synthetase |  |
| *ybtS* | RS21085 | 4.23 | Salicylate synthase |  |
| *ybtU* | RS21120 | 6.30 | Thiazolinyl imide reductase |  |
| *mrkA* | RS02840 | 8.42 | Type 3 fimbria major subunit MrkA | Type 3 fimbria attachment and biofilm<sup>**E**</sup> |
| *mrkB* | RS02835 | 12.64 | Molecular chaperone |  |
| *mrkC* | RS02830 | 10.63 | Fimbria/pilus outer membrane usher protein |  |
| *mrkD* | RS02825 | 10.09 | Fimbrial protein |  |
| *mrkF* | RS02820 | 16.72 | Fimbrial protein |  |
| *mrkH* | RS02805 | 7.09 | PilZ domain-containing protein |  |
| *mrkI* | RS02810 | 6.38 | Helix-turn-helix transcriptional regulator |  |
| *mrkJ* | RS02815 | 6.44 | EAL domain-containing protein |  |

<sup>**A**</sup>,<sup>**B**</sup> <sup>and</sup> <sup>**C**</sup>Gene names, locus tags (abbreviated from NDX50\_RSXXXX) and gene products as annotated for *K. oxytoca* AHC-6 (GenBank Accession No. NZ\_CP098757.1). <sup>**D**</sup>Fold changes in transcript levels of *K. oxytoca* AHC-6 WT 6 h compared to 3 h in LBG medium. <sup>**E**</sup>Predicted function according to Long *et al*. (92).

### TilR is a distinct MarR-family regulator in KoSC

The broad regulatory scope of TilR prompted us to compare its evolutionary and structural features to other KoSC MarR-family proteins. Six additional MarRs (herein designated MarR2-7) were identified in the *K. oxytoca* AHC-6 genome; all showed low pairwise sequence identity with TilR (<40%, **Figure S8A**) and lacked its distinctive protruding α1/α1′ helices (**Figure S8B**). Phylogenetic analysis across all surveyed clinical KoSC isolates placed TilR in a distinct clade from MarR2-7 (**Figure S9**).

To further assess the evolutionary and structural distinctiveness of TilR, we used sequence and structural similarity metrics to compare the seven AHC-6 MarR-family proteins with the 95 structurally characterized MarR-family proteins in the PDB. This analysis identified solved structures closely related to MarR2-7 but not TilR (**Figure S10**). The closest relatives of MarR2-7 included HosA from *E. coli* (8AGA) for MarR2 (93); PecS from *Pectobacterium atrosepticum* (8YLI) for MarR3 (94, 95); RovA from *Yersinia pseudotuberculosis* (4AHI) for MarR4 (96); the uncharacterized MarR-family regulator PA3341 from *Pseudomonas aeruginosa* (2FBH) for MarR5; canonical MarR from *E. coli* (3VOD) for MarR6 (60); and SlyA from *Salmonella enterica* (3DEU) for MarR7 (38) (**Figure S10**). Some MarRs, such as OhRs from *Xanthomonas campestris* and *Bacillus subtilis,* as well as MgrA from *Staphylococcus aureus*, contain redox-responsive cysteine residues that regulate DNA-binding activity (58, 97), but these did not align with any of the three cysteines in TilR (**Figures S11A** and **B**). Consistent with a distinct placement of TilR among MarR-family regulators, BlastP analysis identified the closest TilR sequence homolog in the phylogenetically distant *Xenorhabdus* spp. (65% sequence identity, **Figures S12A and B**), a TM-and TV-producing symbiont of nematodes (98–100). Interestingly, all examined *K. oxytoca* isolates that lacked the TM biosynthesis genes (18, 49) contained *tilR*, *npsC* and *mfsX-uvrX* (**Figures S9** and **S13**), suggesting that these genes represent an ancestral regulatory and transport module that existed prior to acquisition of the complete TM biosynthetic locus. This organization is more consistent with canonical MarR-family loci (42, 59) and may therefore reflect the ancestral state from which the TM-associated *til* locus evolved.

We next examined the genomic context of MarR2-7 to determine whether genomic organization further distinguishes the *tilR* locus from other KoSC MarR-family regulators (**Figure 7**). Similar to *tilR*, all six paralogs were located adjacent to one or more genes encoding multidrug efflux pumps or transporters. However, the *tilR* locus exhibited a distinct organization, with *tilR* located considerably downstream of its associated efflux gene, *mfsX*.

**Figure 7.**
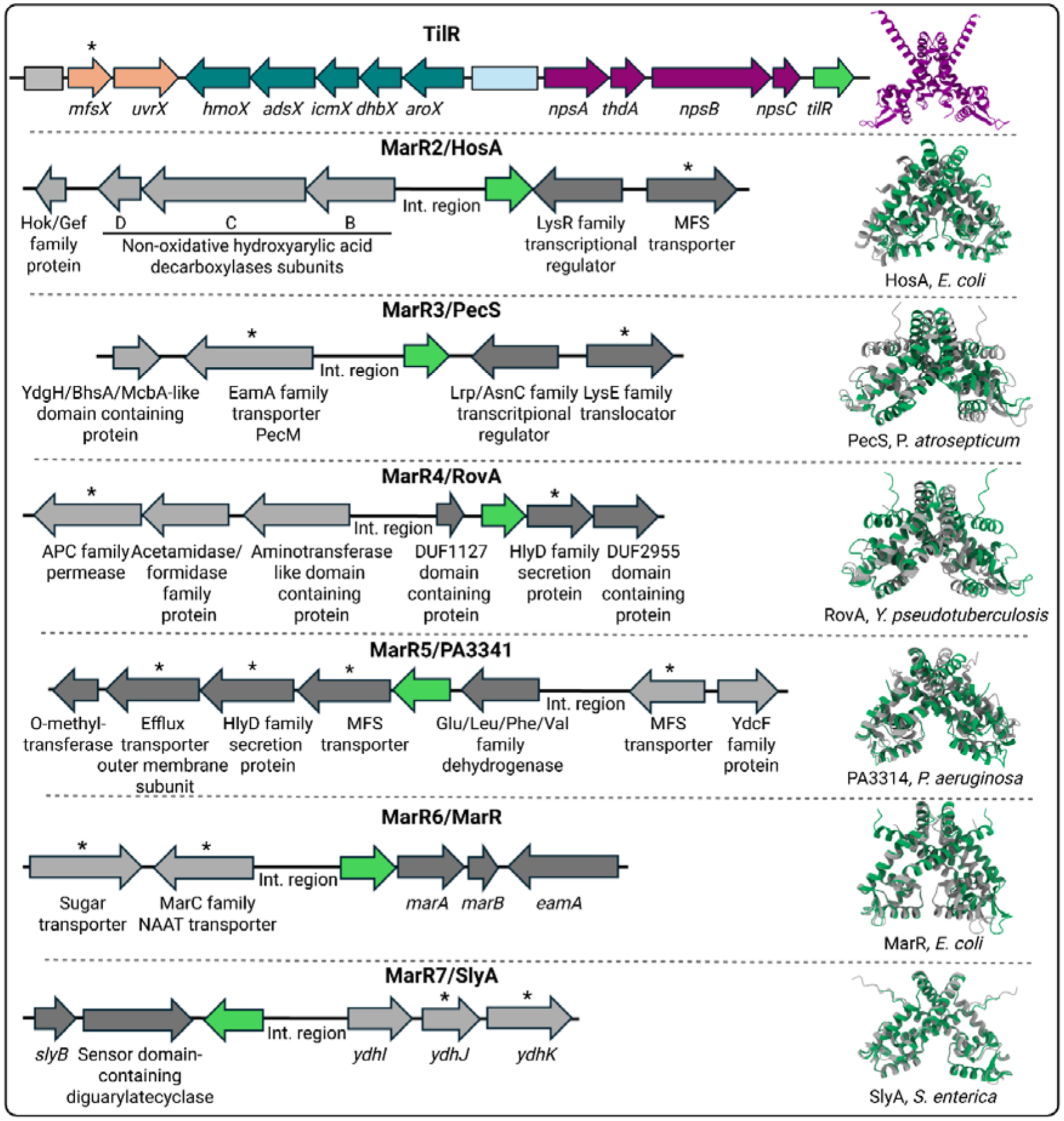
Genomic architecture of loci encoding MarR paralogs in *K. oxytoca* AHC-6. Gene names and gene products are annotated in *K. oxytoca* AHC-6 (GenBank Accession No. NZ_CP098757.1) or in *Salmonella enterica* (genes *ydhI*, *ydhJ* and *ydhK*). Genes encoding putative transporters are marked with asterisks. The genetic loci shown span the following regions: TilR (RS24900-RS24840), MarR2 (RS01960-RS01990), MarR3 (RS02855-RS02875), MarR4 (RS10915-RS10945), MarR5 (RS20120-RS20155), MarR6 (RS21825-RS21850) and MarR7 (RS22625-RS22650). Locus tags were abbreviated from NDX50_RSXXXX to RSXXXX. *K. oxytoca* AHC-6 TilR and MarR2-7 structures (in green, AF3 predicted) are superimposed with structures of well characterized MarR homologs in diverse species (in grey) identified in **Figure S10**. PDB codes for the MarR2-7 homologs are as follows: HosA (8AGA), PecS (8YLI), RovA (4AIH), PA3341 (2FBH), MarR (3VOD) and SlyA (3DEU).

Elevated expression levels are linked to the expanded regulatory repertoires of certain pleiotropic MarR-family regulators (38, 39). We therefore compared the expression of *tilR* with that of MarR2-7. RNA-seq revealed that *tilR* was the most highly expressed MarR-family paralog in WT cells at both time points examined (**Table 2**). Interestingly, *tilR* transcript levels increased approximately 10-fold between 3 and 6 h, coinciding with increased production of TM (**Figure 3**), suggesting that TilR does indeed exert an auto-repressive effect. The other MarRs, in contrast, exhibited little or no change in expression during the transition from early to late growth (**Table 2**). Collectively, these observations identify TilR as a uniquely adapted MarR-family regulator within KoSC, distinguished by its evolutionary divergence, structural distinctiveness, atypical genomic organization, and markedly higher expression than other KoSC MarR-family regulators.

**Table 2.** Transcript levels of TilR and MarR2-7 in *K. oxytoca* AHC-6 WT at 3 and 6 h.

| **MarR paralog** | **Locus tag<sup>A</sup>** | **Homologs<sup>B</sup>** | **3 h<sup>C</sup>** | **6 h<sup>C</sup>** |
|---|---|---|---|---|
| TilR | RS24840 | - | 3,518.76 | 35,972.03 |
| MarR2 | RS01980 | HosA | 53.98 | 75.53 |
| MarR3 | RS02865 | PecS | 9.49 | 13.41 |
| MarR4 | RS10935 | RovA | 26.50 | 42.65 |
| MarR5 | RS20140 | PA3341 | 7.50 | 12.11 |
| MarR6 | RS21835 | MarR | 58.74 | 174.60 |
| MarR7 | RS22635 | SlyA | 1,093.34 | 1,094.01 |

**<sup>A</sup>** The locus tags are abbreviated from NDX50\_RSXXXX to RSXXXX. **<sup>B</sup>** Homologs identified for MarR2-7. <sup>C</sup> Normalized DESeq2 counts of *K. oxytoca* AHC-6 WT at 3 vs 6 h are displayed.

### KpSC contains orthologs for Ko MarR2-7 but not TilR

KpSC and KoSC are principal ecological rivals within the mammalian gut (18, 49, 55). The ability of *K. oxytoca* to outcompete *K. pneumoniae* is thought to involve both toxin-dependent (24) and toxin-independent mechanisms, including enhanced utilization of diverse carbon sources (49, 101). Notably, we identified orthologs of MarR2-7, but not TilR, across representative human KpSC isolates spanning multiple phylogroups (**Figure S14**). Of the three KpSC MarRs that have been functionally characterized, PecS (54) is homologous to MarR3, and KvrA (53) corresponds to MarR7, while KvrB (53) has no close ortholog within KoSC. Comparison of genomic neighborhoods revealed that the MarR2, MarR6, and MarR7/KvrA loci were conserved between *K. oxytoca* AHC-6 and representative *K. pneumoniae* strains (**Figure 7**), while those surrounding MarR3/PecS, MarR4, and MarR5 exhibited lineage-specific divergence (**Figure S15**).

## DISCUSSION

Members of the KoSC, ubiquitous colonizers of the GI tract, are established causes of AAHC in children and adults (15, 16) and have emerged as important contributors to NEC in premature infants (17–19). Central to their pathogenic potential is the production of the enterotoxins TM and TV (21, 22, 24, 102). TM also acts as a mutagen in host cells (25), highlighting the potential long-term oncogenic consequences of KoSC colonization. In this study, we demonstrate, using the clinical isolate *K. oxytoca* AHC-6, that the MarR-like transcription factor TilR, the most highly expressed of the seven encoded MarR-family regulators, represses TM biosynthesis while functioning as a global regulator of efflux and transport systems, nutrient acquisition, stress responses, and central metabolism. Compared with canonical MarRs (42, 44), TilR exhibits a distinct overall structure and genetic locus architecture and is absent from members of KpSC, its closely related rival in the GI tract (18, 49, 55). Together, these findings support a model (**Figure 8**) in which TilR acts as an integrative regulator that couples environmental sensing to toxin synthesis and adaptive transcriptional reprogramming, thereby promoting bacterial fitness and pathogenicity in the gut.

**Figure 8.**
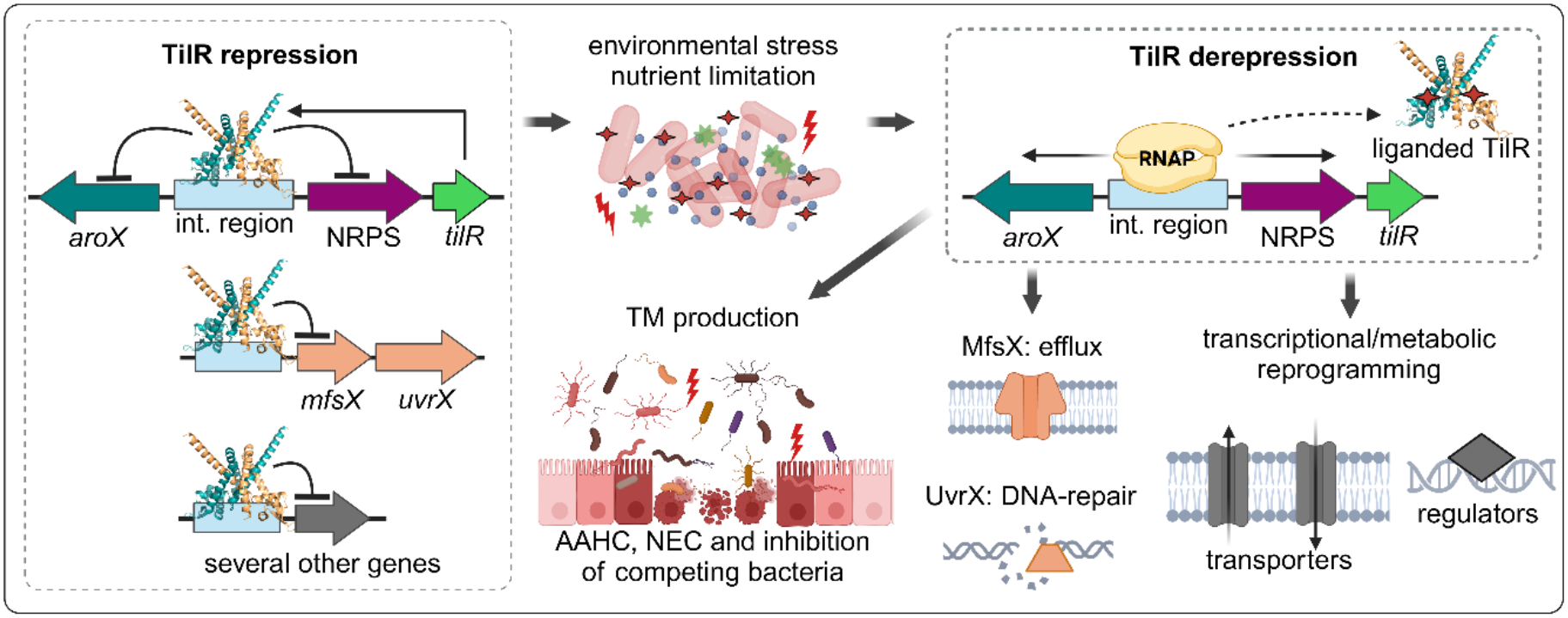
Proposed model for TilR function. During early growth phase, when nutrients are abundant, TilR binds to the intergenic region and directly represses transcription of the *aroX-* and NRPS-operons, *mfsX*-*uvrX*, and additional target genes. During the late growth phase, nutrient limitation triggers the release of environmental stress signals and putative ligands for TilR. Ligand-bound TilR dissociates from its DNA binding site, relieving repression of TilR-regulated loci, including the TM biosynthetic genes. Expression of the TM biosynthetic genes leads to production of TM, resulting in intestinal injury and enabling Ko to out-compete other intestinal commensals. De-repression of TilR-regulated loci also contributes to broader transcriptional and metabolic reprogramming through both direct and indirect regulatory effects.

Canonical MarRs rely on a conserved wHTH motif for DNA binding, with both the α4 helix and wing region contributing to DNA recognition (103, 104). As with other MarRs, TilR residues within α4 and the wing are predicted to mediate DNA interactions, although the precise determinants of binding remain unexplored. TilR diverges from its KoSC paralogs and canonical MarRs, however, through its extended α1 and α’1 helices, structural elements that in some MarRs participate in ligand binding (105). Despite extensive phylogenetic and structural comparisons across the MarR family, no functionally characterized TilR homolog could be identified. This presumed functional specialization becomes evident under nutrient-limiting conditions, as stationary phase signals produced during *in vitro* cultivation appear to relieve TilR-mediated repression, likely through ligand-dependent modulation of DNA-binding activity. Identifying ligands that relieve TilR-mediated repression will be critical for understanding the *in vivo* luminal cues that trigger cytotoxicity. Because biosynthesis of TM intersects with aromatic metabolism (**Figure 1**), shikimate-and chorismate-derived metabolites represent strong candidate signals, supported by parallels with SlyA in *Salmonella enterica* (48). In KoSC, fermentable carbohydrates enhance toxin production (26, 29, 50), whereas indole – a tryptophan-derived bacterial metabolite associated with carbon source depletion and amino acid sufficiency – represses toxin production (26). These observations suggest that luminal conditions favoring indole production may limit the generation or availability of TilR ligands. Given the role of TilR as a global regulator, it is tempting to speculate that multiple endogenous ligands modulate its binding activity in a physiological, context-dependent manner.

TilR represses TM biosynthesis genes by binding a 20 bp DNA motif composed of two outer 6 bp inverted repeats within the *aroX*-NRPS intergenic region, a proven regulatory hotspot (33–36). Supporting this motif as the TilR binding site, loss of a single nucleotide within one of these 6 bp inverted repeats was sufficient to abolish TilR binding. Use of RNA-seq to delineate a transcriptionally silent interval within the *aroX*-NRPS intergenic region improved the accuracy of promoter prediction, enabling the identification of candidate promoters for both *aroX* and *npsA* outside the TilR-binding motif. This refined definition of the intergenic region may explain discrepancies with previously reported promoter annotations (33–36), including those from the recent TagR/TilR study, which proposed an alternative promoter architecture and linked toxin repression primarily to the NRPS operon (33). In contrast, our RNA-seq-guided analysis demonstrated that TilR represses transcription of both the *aroX* and NRPS operons, consistent with bidirectional regulation from a central intergenic binding site.

The arrangement of known regulatory elements within this region also provides insight into the mechanism of repression. Four transcriptional activators – CRP, LRP, Fis, and IHF – bind within the *aroX*-NRPS intergenic region (34–36). Because the binding sites of CRP, Fis, and IHF do not overlap the TilR motif, repression is unlikely to result from direct competition for these factors. In contrast, the LRP binding site partially overlaps the TilR motif, supporting the possibility that TilR and LRP compete for access to this region. Collectively, these observations suggest that TilR-mediated repression may involve modulation of local DNA architecture and/or interference with activator-dependent transcription. In addition to regulating TM biosynthesis genes, TilR also appears to repress its own expression, a hallmark of many MarR-family regulators (42, 59). Consistent with this model, *tilR* expression increases markedly during later growth, in parallel with TM biosynthesis, when TilR is predicted to be ligand-bound and dissociated from DNA. This accumulation of TilR may prime the system for rapid re-establishment of repression once ligand concentrations decline. Given the complex regulatory landscape of the *aroX*-NRPS intergenic region, additional studies will be required to define the precise molecular mechanisms involved.

Most MarR-family regulators typically regulate divergently oriented efflux genes separated by a shared intergenic region (**Figure 7**) (42, 59). In contrast, *tilR* is located several genes downstream of the TilR-regulated *mfsX-uvrX* operon (**Figure 1**), which encodes a putative multidrug efflux pump and DNA repair protein proposed to mitigate TM-mediated autotoxicity and DNA damage (24, 106). The presence of *mfsX* and *uvrX* in toxin-negative *K. oxytoca* strains suggests that regulation of this operon may represent an ancestral function of TilR. One possibility is that TilR-mediated regulation of *mfsX-uvrX* evolved before acquisition of the TM biosynthesis genes, which were integrated subsequently into the TilR regulon. Alternatively, the entire *til* gene cluster, including *tilR*, *mfsX,* and *uvrX*, may have been introduced as a pathogenicity island, most plausibly from *Streptomyces* (21), after which some strains lost the TM biosynthesis genes. The selective pressures driving the loss of TM biosynthesis genes from some KoSC strains remain difficult to reconcile, however, with the presumed fitness benefits of toxin production.

Beyond its unusual genomic organization, TilR also differs functionally from most MarR-family regulators. Although MarRs were originally characterized as transcriptional repressors with narrowly defined regulons, several, including SlyA and PecS, have been shown to function as global regulators (38, 43). Consistent with this broader regulatory role, the TilR regulon extends well beyond the TM biosynthesis genes and the *mfsX–uvrX* operon. *K. oxytoca* encodes seven MarR paralogs, including homologs of SlyA and PecS; however, under the conditions tested herein, TilR was the most highly expressed and the only paralog that exhibited substantial growth phase-dependent induction. Elevated expression has been linked to expansion of regulatory repertoires in other pleiotropic MarR-family regulators (39) and may likewise have contributed to the emergence of TilR as the dominant MarR regulator in KoSC, whereas the contributions of other paralogs may be limited to narrower, context-dependent regulatory roles.

Particularly notable is the TilR-dependent repression of transcriptional regulators associated with carbon metabolism (*pdhR*), oxidative stress and metal homeostasis (MerR-family regulator), cell envelope maintenance (TraR-family regulator), and global metabolic control (AraC-and LysR-family regulators). Among these is an AraC-family regulator related to MarA of the canonical MarRAB system (107). Although the physiological roles of these regulators in KoSC remain to be defined, their predicted functions suggest that TilR exerts some of its effects through downstream regulatory networks. For example, PdhR functions as a sensor of central carbon flux that contributes to virulence regulation in diverse bacterial species (78, 79, 108), raising the possibility that TilR links toxin production to cellular metabolic state through a regulatory hierarchy. Consistent with this possibility, many TilR-regulated genes lack obvious matches to the inferred TilR-binding motif, suggesting that TilR regulates gene expression through a combination of direct and indirect mechanisms. For genes regulated directly by TilR, variation among binding sites may be tolerated at some positions, whereas a limited number of critical nucleotides likely dictate binding specificity. Such flexibility has precedent within the MarR family, as PecS in *Erwinia chrysanthemi* binds a defined consensus motif but also several promoters lacking a clear match (43, 109). Similar combinations of sequence-specific and sequence-permissive DNA recognition occur in some FUR-family regulators (110, 111), nucleoid-associated proteins such as Fis, H-NS, and IHF (112–114), and other DNA-binding proteins (113).

An unexpected finding was that several genes involved in oxidative and nitrosative stress responses, including the *hcp-hcr* operon, required TilR for expression during late growth, when TilR is predicted to be ligand-bound and largely dissociated from DNA. Because TilR-dependent transcriptional regulation is largely absent at this stage, these expression changes are likely indirect and may arise from downstream effects of the broader metabolic dysregulation caused by loss of TilR during earlier growth phases. Defining the mechanistic links between these early regulatory events and late-stage stress-response gene expression will require further investigation.

In the competitive gut ecosystem, coordinated deployment of toxin production, nutrient acquisition, and metabolic reprogramming is likely to confer a substantial fitness advantage unique to KoSC. TilR synchronizes TM production with enhanced uptake of diverse carbon sources, coupling suppression of competing microbes to efficient exploitation of host-derived nutrients. This integrated strategy may enhance colonization resistance against *K. pneumoniae* and other enteric pathogens (49, 50, 101), a particularly relevant outcome given that invasive *K. pneumoniae* infections often originate from intestinal reservoirs (115, 116). The absence of TilR and its regulon in members of the KpSC points to an evolutionarily specialized adaptation that contributes to the ecological success of KoSC. Importantly, TM production coincides with induction of additional TilR-independent virulence-associated factors, indicating that toxin synthesis occurs within a broader host-adaptive physiological state. These functions may enhance competitive fitness by acting in concert with TM-mediated inhibition of competing bacteria, while also contributing to disease pathogenesis. For example, increased adherence could promote localized or sustained toxin delivery to the intestinal epithelium, exacerbating mucosal injury. This coordinated physiological state provides a plausible mechanistic framework for diseases associated with KoSC, such as AAHC and NEC (15–19). Defining toxin-independent functions of TilR will be critical for understanding KoSC ecology, particularly its role in colonization resistance and broader host-microbe interactions.

## MATERIALS and METHODS

### Bacterial strains and plasmids

*K. oxytoca* AHC-6 (GenBank accession no. CP098757.1), isolated from a patient with AAHC, is naturally resistant to β-lactam antibiotics and harbors a kanamycin resistance marker (*aphA*) (21). The AHC-6 Δ*tilR* and plasmids p*tilR*, p*tilR*-HA, and pIPTG-*tilR*-HA were constructed as described below (**Table S1** lists all plasmids used in this study). pAR181 was a kind gift from E. Zechner (University of Graz, Austria). *K. oxytoca* and *E. coli* strains were grown in Luria–Bertani broth (LB) or LB supplemented with 2.5 g/L glucose (LBG) at 37 °C with shaking at 250 rpm; growth was monitored by measuring the OD_600_. Strains harboring plasmids were cultivated in the presence of the appropriate antibiotics (chloramphenicol, 40 µg/mL; spectinomycin, 100 µg/mL; apramycin, 50-70 µg/mL; streptomycin, 100 µg/mL).

### Routine DNA manipulation and cloning

CloneAmp HiFi Polymerase was used for routine PCR amplifications and cloning with the In-Fusion HD Cloning Plus kit (both from TaKaRa Bio USA, Inc.). Oligonucleotides (**Table S2**) were purchased from Sigma-Aldrich (St. Louis, MO). To generate p*tilR*, pAR181 was linearized by inverse PCR with oligonucleotides pAR181-c*tilR*_F and pAR181-c*tilR*_R. The GFP cassette was replaced with the entire *aroX*-NRPS intergenic region amplified from *K. oxytoca* AHC-6 genomic DNA using c*tilR*-pAR_F and c*tilR*-pAR_R. To generate p*tilR*-HA, a HA tag was added to the TilR-coding sequence in *tilR* by inverse PCR of p*tilR* using *tilR*-HA-pAR181_F and *tilR*-HA-pAR181_R. An empty-vector control plasmid (pAR181-empty) was generated using empty_pAR181_F and empty_pAR181_R. The inducible plasmid pIPTG-*tilR*-HA was generated in two steps. First, pJSB275 was modified by inverse PCR using pJSB275-mod_F and pJSB275-mod_R to remove certain vector components. Second, using pJS-*tilR*HA_F and pJS-*tilR*HA_R for inverse PCR, the luciferase gene was replaced by *tilR*-HA amplified from p*tilR*-HA using *tilR*HA_pJS_F and *tilR*HA_pJS_R. PCR fragments were cloned into the linearized plasmids using In-Fusion cloning. Plasmids were transformed into chemically competent *E. coli* Stellar cells (TaKaRa Bio USA, Inc., San Jose, CA) by heat shock, recovered for 1 h at 37 °C, and plated on LB agar supplemented with selective antibiotics. Purified plasmids were transformed into competent *K. oxytoca* AHC-6 cells using the same procedure. Sanger sequencing was performed by Genewiz, Inc. (South Plainfield, NJ) and analyzed using MacVector software (MacVector, Inc., Apex, NC).

### Generation of a *tilR* deletion mutant

A *Klebsiella oxytoca* AHC-6 Δ*tilR* mutant was generated using a modification of the pCasKP-pSGKP clustered regularly interspaced short palindromic repeat (CRISPR)-Cas9/lambda Red recombination system described by Wang *et al.* (117). pCasKP-apr (catalog no. 117231) and pSGKP-spe (catalog no. 117234) were obtained from Addgene (Watertown, MA). The heat-sensitive pCasKP-apr (apramycin resistance) encodes the Cas9 nuclease and the lambda Red recombination genes (*gam*, *bet*, and *exo*). pSGKP-spe (spectinomycin resistance) contains a single artificial chimeric guide RNA (sgRNA) and the *sacB* gene for plasmid curing. The Cas9-generated double-strand DNA break was repaired by the lambda Red recombination system using a short repair fragment. The sgRNA spacer sequence was generated with *tilR*-spacer_F and *tilR*-spacer_R and cloned into pSGKP. The 1-kb DNA repair fragment extending 500 bp upstream and downstream of the *tilR* gene was generated using overlap-extension PCR. The upstream fragment (UP) was amplified from *K. oxytoca* AHC-6 genomic DNA using *tilR*-UP_F and *tilR*-UP_R, while the downstream fragment (DW) was amplified using *tilR*-DW_F and *tilR*-DW_R. 25 ng each of the UP and DW fragments were used in a primer-less PCR for 15 cycles to allow fragment annealing, following which *tilR*-UP_F and *tilR*-DW_R were added and PCR was continued for 35 cycles. The spacer-containing pSGKP plasmid and the repair fragment were transformed into *K. oxytoca* AHC-6 via electroporation; competent cells were pulsed at 1.8 kV and recovered for 90 min at 30 °C and selected on LB agar containing spectinomycin and apramycin. Successful deletion of *tilR* was confirmed by PCR with *tilR*_F and *tilR*_R and sequencing; the pCasKP and pSGKP plasmids were cured according to Wang *et al.* (117).

### Detection of tilimycin by ultra-performance liquid chromatography-tandem mass spectrometry (UPLC-MS/MS)

*K. oxytoca* AHC-6 strains were grown in LBG for 4, 6 and 8 h and pelleted at 8,000 × g for 6 min at RT; supernatants were cleared using 0.2 µm surfactant-free cellulose acetate (SFCA) filters; one mL aliquots were stored in glass vials at-20 °C prior to methanol extraction and measurement of TM. Samples were analyzed at the University of Connecticut Center for Environmental Sciences and Engineering (Storrs, CT) using a Waters Acquity UPLC coupled with an Acquity TQD tandem mass spectrometer (Waters Co., Milford, MA) as previously described (26). A total of 450 µL of test or quality control samples were spiked with 25 µL of ketoprofen-d3 internal standard solution and 25 µL of methanol prior to injection. An Acquity UPLC BEH C18 column (1.7 µm, 2.1 x 50 mm), maintained at 25 °C with a sample injection volume of 10 µL on a 20 µL loop, was utilized for analyte gradient separation. Analyte signal optimization was performed using the Waters IntelliStart, whereas analyte detection, quantification, and statistical analysis were performed using the Waters QuanLynx within MassLynx software v.4.2. Electrospray positive ionization (ESI+) was performed in multiple-reaction monitoring (MRM) MS scan mode.

### Cytotoxicity assay

HT-29 colon cancer cells, obtained from the American Type Culture Collection (ATCC, Manassas, VA, USA; catalog no. HTB-38), were grown in modified McCoy’s 5A (Gibco, Thermo Fisher Scientific, catalog no. 16600082) medium supplemented with 10% fetal bovine serum (catalog no. A5256701) at 37 °C with 5% CO_2_ in 95% humidity. Cells were seeded in 96-well tissue culture plates at a density of 1.5 × 10^4^ cells/100 μL and left to adhere for at least 12 h. Bacterial cultures of *K. oxytoca* AHC-6 WT and Δ*tilR* at 4, 6 and 8 h of growth in LBG were centrifuged at 8000 × g for 6 min at RT. Supernatants were filtered using 0.2 µm SFCA filters and diluted (1:2) in McCoy’s 5A medium. After 48 h of incubation with diluted supernatants, viable HT-29 cells were measured via staining with 3-(4,5-dimethylthiazol-2-yl)-2,5-diphenyltetrazolium bromide (MTT) (Sigma-Aldrich, catalog no. 475989) as previously described (22). A_595_ values were normalized to negative controls – HT-29 cells treated with supernatants of *K. oxytoca* toxin-negative strain ATCC 13182 (ATCC, catalog no. 13182) – set to 100% viability. Results are presented as percent cell death, calculated as 100 − percent viability.

### SDS-PAGE and immunoblotting

*K. oxytoca* Δ*tilR ptilR-*HA (complementation strain) was cultured in LBG for 2, 4, 6, and 8 h, while the pIPTG*-tilR-*HA strain (used for DNA pull-down assays, see below) was grown for 1 h in LB followed by induction with IPTG for 2 h. Cells were adjusted to an OD_600_ of 1, pelleted at 6000 × g for 3 min at RT, washed once with phosphate-buffered saline (PBS), and resuspended in 90 µL PBS. NuPAGE LDS Sample Buffer (4X, Invitrogen, catalog no. NP0008) supplemented with 4% β-mercaptoethanol (BME) was added (30 µL) and incubated at 99 °C for 10 min. 10 µL of whole-cell lysates were loaded onto Any kD™ Mini-PROTEAN® TGX™ precast protein gels (Bio-Rad, catalog no. 4569033). Volumes of whole-cell lysates were adjusted to ensure equal loading. Gels were either stained using the eStain® L1 protein staining system (GenScript) or transferred to 0.2 µm nitrocellulose membranes (Bio-Rad, catalog no. 1620112) for immunoblotting. Membranes were incubated overnight at 4°C with the primary antibody Anti-HA-Biotin, High Affinity (3F10) (Roche, catalog no. 12158167001), diluted 1:5000 in 5% milk in PBS with 0.1% Tween-20 (PBS-T). After washing in PBS-T, membranes were incubated with horseradish peroxidase-conjugated goat anti-rat secondary antibody (Southern Biotech, catalog no. 3030-05) diluted 1:30,000 in 5% milk in PBS-T. Immunoblots were developed using the Amersham^TM^ ECL detection reagent (Cytiva, catalog no. RPN3004) and the ChemiDoc Imaging System (Bio-Rad).

### DNA pull-down to identify the TilR binding site

*K. oxytoca* AHC-6 Δ*tilR* pIPTG-*tilR*-HA was grown in LBG for 1 h and then induced with 100 µM IPTG for 2 h. Cells were pelleted at 6000 × g for 3 min at RT and then resuspended in 1 mL PBS. After sonication for 1 min (10 sec on, 50 sec off) on ice, lysates were spun at 20,000 × g for 10 min at 4 °C and supernatants containing TilR-HA were used for the pull-down assays. The following biotinylated primer pairs (**Table S2**) were used to generate 5′-biotinylated versions of the entire *aroX*-NRPS intergenic region and fragments 1, 2, 3, 4 and 4a (**Figure 4**): 1, KoIn-btn_F and KoIn_R; 2, KoIn-btn_F and KoIn255_R; 3, KoIn191-btn_F and KoIn_R; and 4, KoIn100-btn_F and KoIn100_R. For fragment 4a, inverse PCR of p*tilR* using MutBin_F and MutBin_R was performed to mutate the TilR binding site, and the 100 bp fragment was amplified using KoIn100-btn_F and KoIn100_R. Streptavidin magnetic beads (Dynabeads^TM^ M-280, Invitrogen, catalog no. 11205D) were used to bind biotinylated DNA (per Dynabeads manual). 10 µL beads were washed three times with 1 mL B&W buffer followed by incubation with the biotinylated DNA (4 µg/200 µL) for 45 min at RT; beads were washed three times with PBS to remove unbound DNA. Bacterial supernatants containing TilR-HA per above were incubated with DNA bound to beads for 2 h 30 min with rotation at RT in PBS. The supernatants were collected and beads washed three times with 1 mL PBS to remove unbound protein and resuspended in 50 µL PBS. 10 µL of lysate, beads and supernatant were mixed with 2 µL NuPAGE LDS Sample Buffer (4X, Invitrogen, catalog no. NP0008) supplemented with 4% BME, incubated at 99 °C for 10 min and 10 µL of each sample were loaded onto the gel. TilR-HA was detected by immunoblotting using Anti-HA-Biotin, High Affinity (3F10) (Roche, catalog no. 12158167001) diluted 1:5000 in 5% milk in PBS-T with membranes incubated overnight at 4 °C.

### RNA isolation, library preparation, and RNA-seq

Total RNA was isolated from *K. oxytoca* AHC-6 WT and Δ*tilR* strains at 3 and 6 h during growth in LBG. Four biological replicates were collected per strain at each time point. RNA was extracted with TRIzol (Thermo Fisher Scientific), treated twice with TURBO DNA-free DNase (Thermo Fisher Scientific), and purified after each DNase treatment using RNeasy columns (Qiagen) according to the RNA Cleanup protocol, as previously described (118, 119). RNA integrity was assessed using a TapeStation 4200 (Agilent Technologies) prior to library preparation. Library construction was performed with the Illumina Stranded Total RNA prep, Ligation kit with Ribo-Zero kit (Illumina, San Diego, CA). Libraries were validated for fragment size distribution (TapeStation 4200 D1000 high-sensitivity assay), quantified using the Qubit dsDNA HS Assay Kit (Thermo Fisher Scientific), and normalized prior to sequencing. High-throughput sequencing was carried out on an Illumina NovaSeq 6000 platform. Reads were quality-trimmed with Sickle (v1.33, https://github.com/najoshi/sickle) and mapped to the *K. oxytoca* AHC-6 reference genome (GenBank accession no. CP098757.1) using EDGE-pro (v1.1.3) (120) with custom FASTA, protein translation table (ptt), and ribosomal/transfer RNA (rnt) annotation files. Pseudogenes and coding sequences <60 amino acids were excluded. Differential gene expression between strains and time points was determined using DESeq2 (v1.26.0), with significantly regulated genes defined as ≥3-fold change and Benjamini–Hochberg–adjusted q < 0.05. Principal component analysis (PCA) was performed in R using variance-stabilized counts generated with DESeq2 (blind = TRUE), based on the 500 most variable genes, and visualized using ggplot2.

### Promoter predictions using transcriptomic data

Strand-specific RNA-seq data were further analyzed to resolve transcriptional organization within the *aroX*-NRPS intergenic region. Quality-trimmed reads were re-aligned to the *K. oxytoca* AHC-6 reference genome (GenBank accession no. CP098757.1) using Bowtie2 (v2.5.0) (121, 122) with default parameters. Alignment files were processed and sorted using SAMtools (v1.20) (123). Directional read coverage was visualized in Integrative Genomics Viewer (IGV, v2.19.7) (124) to distinguish transcription originating from the minus strand (*aroX*-oriented) and plus strand (*npsA*-, *npsC*-, and *tilR*-oriented). Strand-specific coverage profiles were used to identify regions lacking detectable transcription and to infer putative transcription start sites (TSS) based on the boundaries of abrupt read accumulation. Promoter elements within the *aroX*-NRPS intergenic region and upstream of *tilR* were predicted using Neural Network Promoter Prediction (63). Predictions for the *aroX* and *npsA* promoters were guided by the inferred TSS positions and constrained to the transcriptionally silent sequence. The spatial relationship between predicted promoter elements and the TilR binding site was assessed to infer potential mechanisms of transcriptional regulation.

### Structural modeling

TilR structures were generated using AlphaFold3 (125) in the presence and absence of random sequence duplex DNA. Structures of the MarR-like transcription factors MarR2–7 from *K. oxytoca* AHC-6 and the TilR homolog from *Xenorhabdus eapokensis* were also modeled using AF3. Predicted structures were compared with experimentally determined MarR-family protein structures retrieved from the RCSB Protein Data Bank (PDB). Structural superpositions and figure rendering were performed with the PyMOL Molecular Graphics System (v2.3.2; Schrödinger, LLC, New York, NY) and ChimeraX (126). Electrostatic surface potential was calculated using APBS and visualized with PyMOL.

### Sequence analysis, comparative genomics, and phylogenetics

Protein sequences of TilR and MarR2–7 from *K. oxytoca* AHC-6 were used to identify homologs in representative KoSC and KpSC genomes and other bacterial species using BLASTP. Genes annotated as *marR* or *slyA* in NCBI were included in the analysis. Protein domains and Gene Ontology (GO) annotations were assigned using InterProScan (127). Virulence-associated homologs were identified by comparison with the *K. pneumoniae* virulence-factor dataset described by Long *et al*. (92) using InterPro. Genomic neighborhoods of the identified MarR-family proteins were compared using publicly available genome annotations. Experimentally determined MarR-family structures were identified by searching the RCSB Protein Data Bank (PDB). Protein sequences were aligned using Clustal Omega (128) or MAFFT Multiple Sequence Alignment Software Version 7 (129), and alignments were visualized using ESPript 3.0 (ENDscript; https://endscript.ibcp.fr) (130). Phylogenetic analyses were performed using PhyML 3.0 (131), and the resulting trees were visualized using Interactive Tree of Life (iTOL) (132). Inverted repeats within DNA sequences were identified using EMBOSS Palindrome (62). Putative TilR-binding sites were identified using FIMO, and consensus binding motifs were generated using MEME (MEME Suite).

### Statistics

Statistical analyses were performed using GraphPad Prism (v11.0.0), with statistical tests specified in the corresponding figure legends. A P-value less than 0.05 was considered statistically significant. Illustrations were prepared using BioRender.com.

## Data availability

RNA-seq raw data are deposited in NCBI SRA under accession PRJNA1440479. The *Klebsiella oxytoca* AHC-6 genome is available in GenBank under accession no. CP098757.1.

## Author contributions

A.P.M., K.U., J.D.R., M.J.C., and A.A.G. designed and directed experiments; K.U. and A.A.G. performed experiments; K.U., A.P.M., J.D.R., M.J.C., A.A.G., P.B., A.A.P., and Y.Z. analyzed and interpreted data; A.P.M., K.U., J.D.R., M.J.C. and A.A.G. wrote the manuscript.

## Acknowledgements

We thank Ferric C. Fang and Ellen L. Zechner for helpful discussions and critical review of the manuscript. We also thank Dr. Zechner for providing the *K. oxytoca* AHC-6 strain and pAR181 plasmid.

