## Supplemental Material for "The MarR-family regulator TilR links biosynthesis of enterotoxin tilimycin with physiologic adaptation in *Klebsiella oxytoca*"

Adam P. Matson, MD, MSc

Department of Pediatrics

UConn Health

263 Farmington Avenue

Farmington, CT 06030

**Table S1.** Plasmids used in this study.

| **Name** | **Description** | **Use** | **AB resistance** | **Reference** |
| --- | --- | --- | --- | --- |
| pCasKP-apr | Cas9 protein and lambda Red recombination genes | Generation of Δ*tilR* | Apramycin | (1) |
| pSGKP-spe | sgRNA and *sacB* | Generation of Δ*tilR* | Spectinomycin | (1) |
| pAR181 | pAR181 with Pr*_npsA_*-*gfpmut3b* cloned in SacI/SphI sites | Backbone vector for complementation of Δ*tilR* | Chloramphenicol | E. Zechner  lab |
| p*tilR* | pAR181 encoding the *aroX*-NRPS intergenic region and *tilR* | Complementation of Δ*tilR* with expression of TilR driven by the *aroX*-NRPS intergenic region | Chloramphenicol | This study |
| p*tilR*-HA | pAR181 encoding the *aroX*-NRPS intergenic region and *tilR-*HA (C-terminal) | Complementation of Δ*tilR* with expression of TilR-HA driven by the *aroX*-NRPS intergenic region | Chloramphenicol | This study |
| pAR181-empty | Empty vector | Control plasmid | Chloramphenicol | This study |
| pJSB275 | pJSB104 lacking the NdeI site in the resistance marker | Backbone vector for inducible TilR-HA construct | Spectinomycin/ streptomycin | (2) |
| pJSB275-mod | Partially modified pJSB275 | Intermediate construct for pIPTG-*tilR*-HA | Spectinomycin/ streptomycin | This study |
| pIPTG-*tilR*-HA | pJSB275 encoding IPTG-inducible *tilR*-HA | IPTG-inducible expression of TilR-HA | Spectinomycin/  streptomycin | This study |

**Table S2.** Oligonucleotide primers used in this study.

| **Name** | **Sequence 5’-3’** |
| --- | --- |
| **Deletion of *tilR*** (***** used for sequencing) | |
| *tilR*-spacer_F | **TAGT**TGAATTCTGCAGAGTCCAGC (restriction site in bold) |
| *tilR*-spacer_R | **AAAC**GCTGGACTCTGCAGAATTCA (restriction site in bold) |
| *tilR*-UP_F | GATAATACAATGGTTCCAGCAC* |
| *tilR*-UP_R | CGCTTCATCACCATAAACTTTAACATTCTCCTGATGCCG |
| *tilR*-DW_F | CGGCATCAGGAGAATGTTAAAGTTTATGGTGATGAAGCG |
| *tilR*-DW_R | AAGGAAGAGAACAAAATGCAC* |
| *tilR*_F | TATTTCCAGCGGCAGAAAG |
| *tilR*_R | TCGATCATTATCAGCAGGGG |
| **Complementation of Δ*tilR*** | |
| pAR181-c*tilR*_F | GCTTAATTAGCTGAGCTTGGACTCC |
| pAR181-c*tilR*_R | CACTCTCTCCTGGAGAATTAGG |
| c*tilR*-pAR_F | CTCCAGGAGAGAGTGATGGAAATGAATAAAAAAGTGGCT |
| c*tilR*-pAR_R | CTCAGCTAATTAAGCTTAATTTTCACTGGTATCTTCGAGT |
| **C-terminal HA-tagged TilR** (add HA tag to p*tilR*) | |
| *tilR*-HA-pAR181_F | TAAATCGATAAGCTTTAATGCGGT |
| *tilR*-HA-pAR181_R | AAGCTTATCGATTTAGGCATAATCCGGCACATCATACGGATAATTTTCACTGGTATCTTCGAGTTT |
| **Generation of an empty pAR181** | |
| empty_pAR181_F | TCGAGCTCGCTTAATTAGCTGAGCTTGGACTCC |
| empty_pAR181_R | ATTAAGCGAGCTCGAATTCGCGCGG |
| **IPTG-inducible TilR-HA** | |
| pJSB275-mod_F | TATGGAAATCTAGTCTAGTGCCGGCCG |
| pJSB275-mod_R | GACTAGATTTCCATAGGCTCCGCCCC |
| pJS-*tilR*HA_F | TAAAAGCTTGAAGATAGAGAGAGAA |
| pJS-*tilR*HA_R | ATGTAATTTCTCCTCTTTAATGAAT |
| *tilR*HA_pJS_F | GAGGAGAAATTACATATGGAAATGAATAAAAAAGTGGCT |
| *tilR*HA_pJS_R | ATCTTCAAGCTTTTATTAGGCATAATCCGGCAC |
| **Biotinylation of full and truncated aroX-NRPS intergenic region fragments** | |
| KoIn-btn_F (448 bp) | btn-TTCTAATCTCTCACTCGAAATTTAACAGG (5′ biotinylated) |
| KoIn_R | CTGGAGAATTAGGAACGTTCACA |
| KoIn255_R | GATTTCCAAAAAAAAATCATATAGC |
| KoIn191-btn_F | btn-ATTGACTGGGGATGAGGTTC (5′ biotinylated) |
| KoIn100-btn_F | btn-TGGTTTTTTATTTTTTGCTATATG (5′ biotinylated) |
| KoIn100_R | AAGCATAACTAAGCGTTGC |
| **Mutation of TilR binding site** | |
| MutBin_F | CGCGTTTTTTGGCGCGACAATTGACTGGGGATGAGGT |
| MutBin_R | GCGCCAAAAAACGCGACTATAGCAAAAAATAAAAAACCAACA |

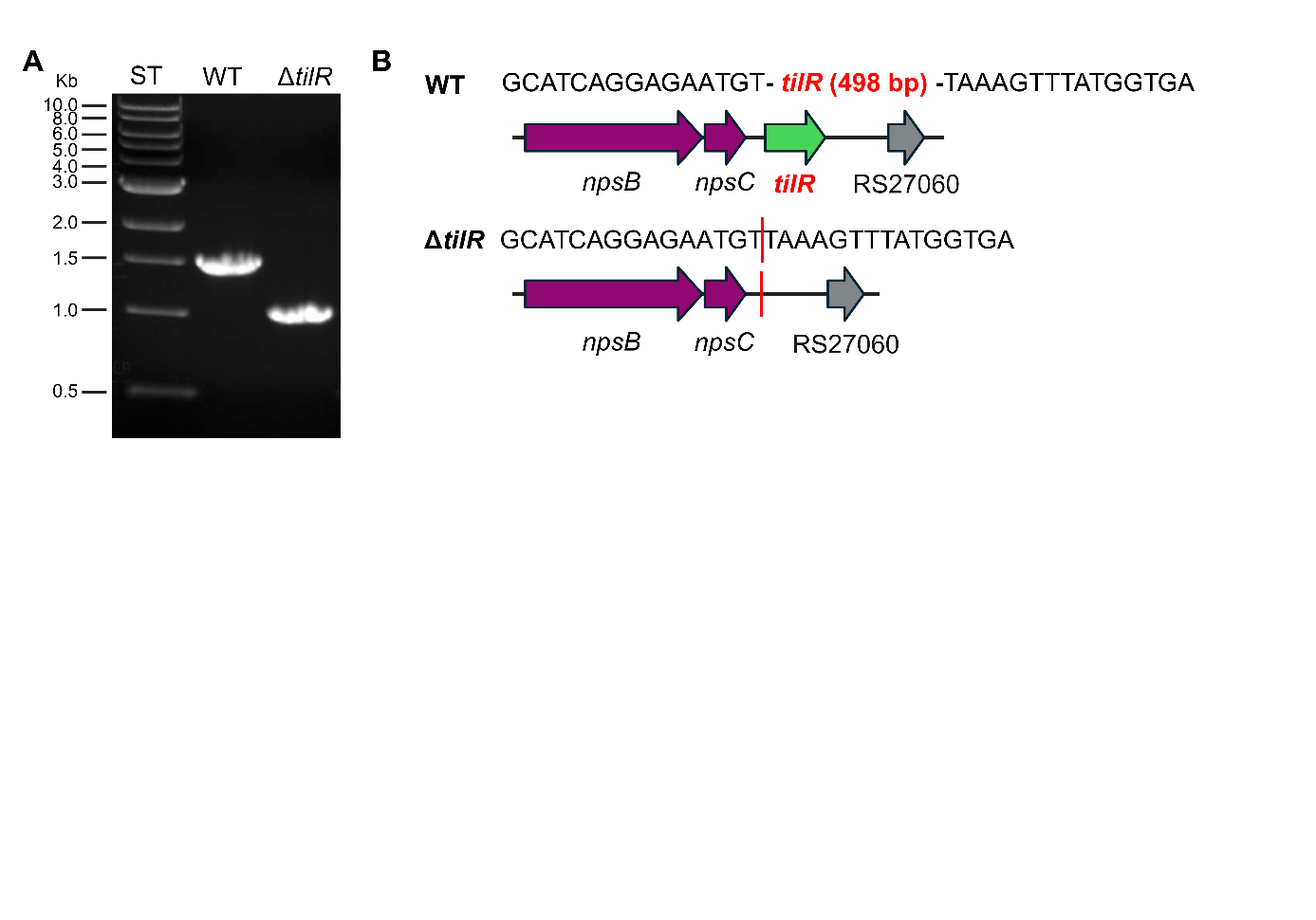
**Figure S1. DNA sequencing and PCR confirm deletion of *tilR.*** (**A**) Size difference of PCR fragments amplified with primers flanking *tilR* (1542 bp for WT and 1044 bp for Δ*tilR*) verifies deletion of *tilR*. (**B**) Sanger sequencing of PCR fragments amplified from WT and the Δ*tilR* strain. Maps show gene names and locus tags abbreviated from NDX50_RSXXXX to RSXXXX, as annotated in *K. oxytoca* AHC-6.

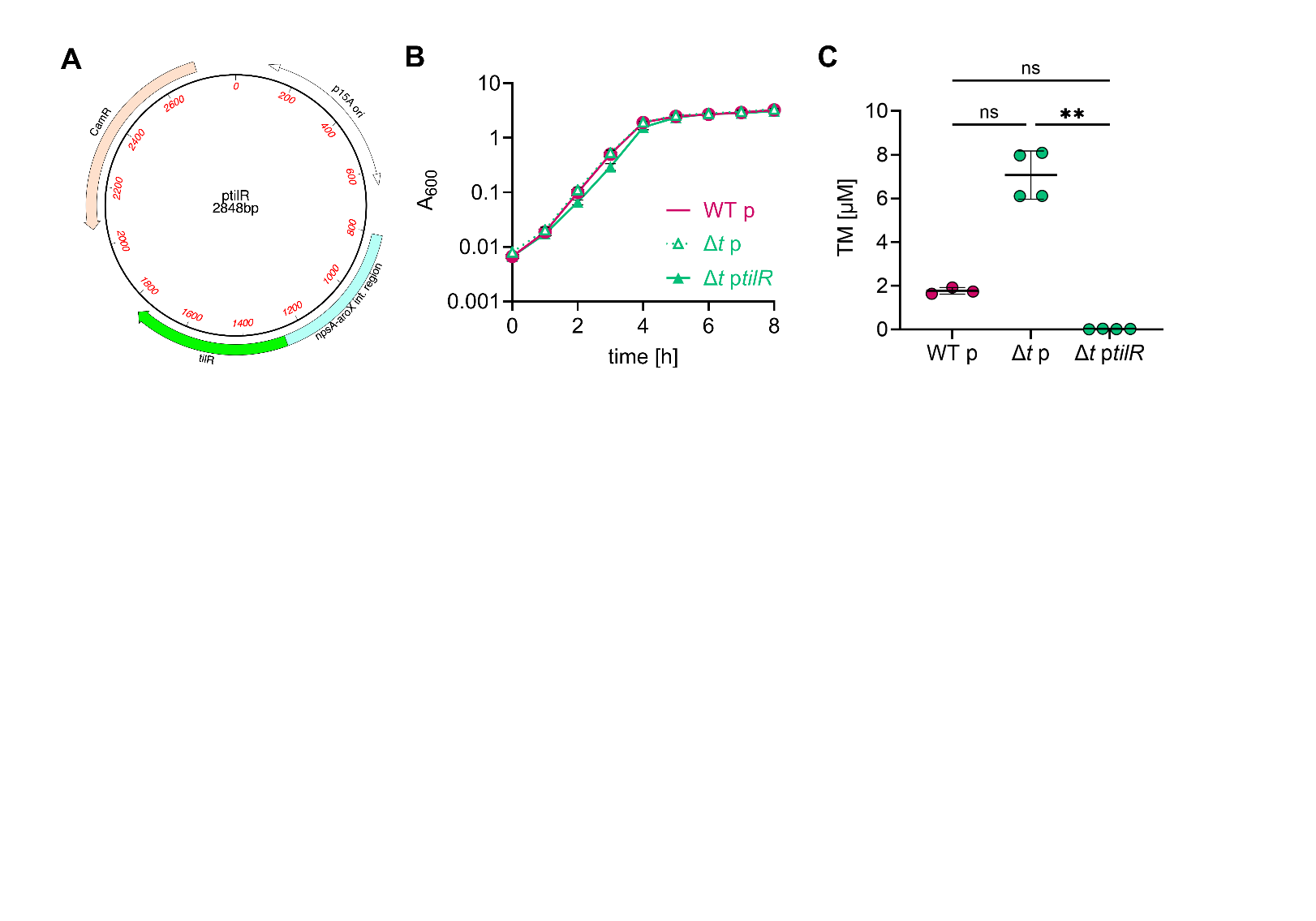

**Figure S2.** **Complementation of the Δ*tilR* mutant restores repression of TM.** (**A**) Plasmid map of p*tilR,* a pAR181 derivative harboring *tilR* downstream of the *aroX-*NRPS intergenic region. (**B**) Growth curves of *K. oxytoca* AHC-6 WT (WT p) and Δ*tilR* (Δ*t* p), both harboring the empty pAR181, and the complementation strain Δ*tilR* p*tilR* (Δ*t* p*tilR)* (mean ± SD, n=3). (**C**) TM concentrations in supernatants following growth for 8 h in LBG (mean ± SD, n=3-4). Statistical analysis was performed using the Kruskal-Wallis test (**P ≤ 0.01).

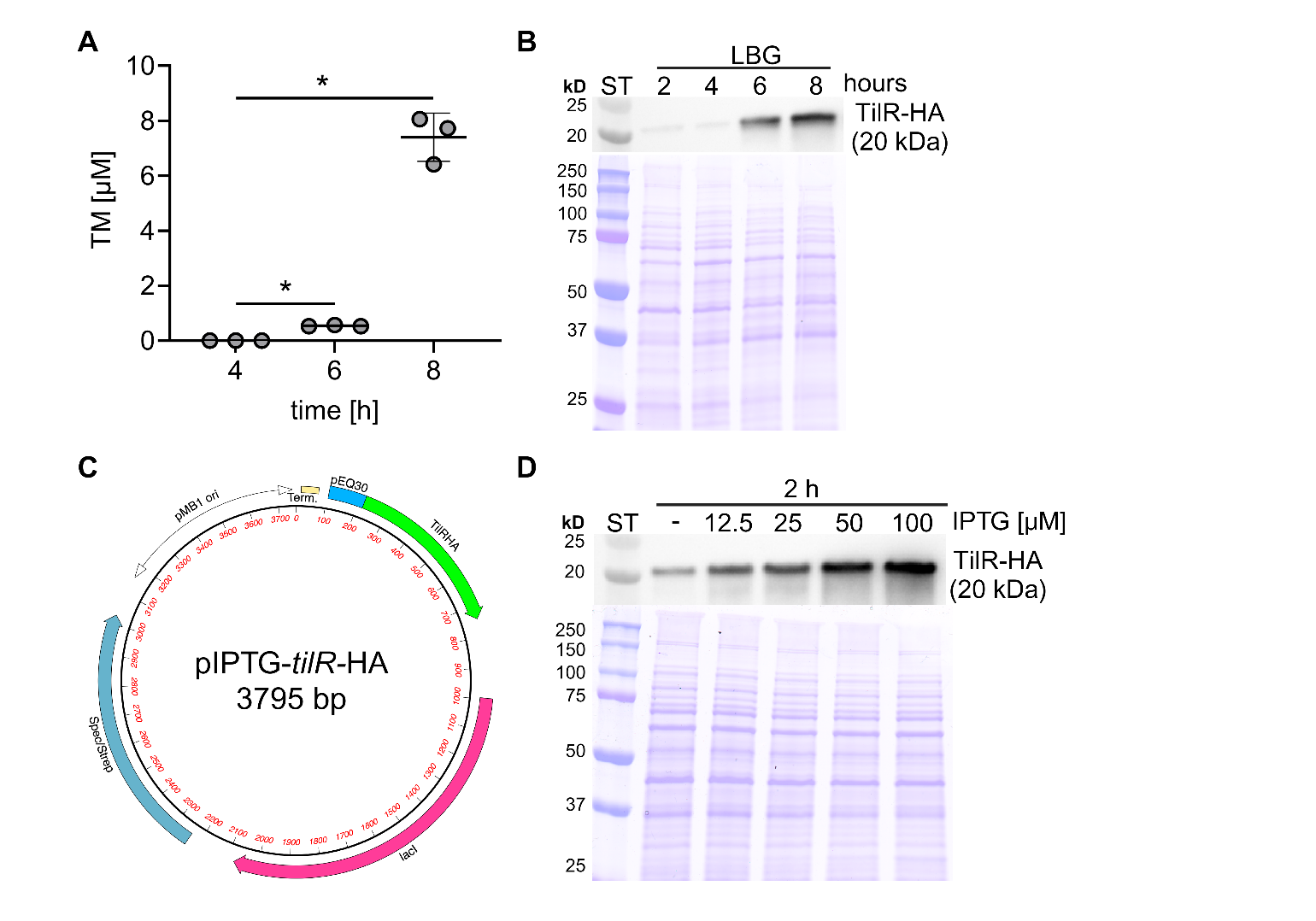
**Figure S3. HA-tagged TilR complements Δ*tilR* in *K. oxytoca* AHC-6.** (**A**) TM levels in supernatants of cultures expressing TilR-HA at 4, 6 and 8 h of growth in LBG (mean ± SD, n=3). Statistical analysis was performed using one-way RM ANOVA followed by Sidak’s multiple comparison test (**P* ≤ 0.05). (**B**) Growth-dependent expression of HA-tagged TilR in *K. oxytoca* AHC-6 Δ*tilR* p*tilR*-HA at 2, 4, 6 and 8 h of growth in LBG (Coomassie blue-stained gel shows equal loading). (**C, D**) To achieve higher expression levels of TilR-HA, the encoding gene was moved to pJSB275, containing an IPTG-inducible promoter, creating pIPTG-*tilR*-HA.

**Table S3.** Fold changes in transcript levels for genes within the *til* gene cluster in *K. oxytoca* AHC-6 WT vs Δ*tilR* at 3 h and 6 h.

| **Gene name^A^** | **Locus tag^B^** | **WT vs Δ*tilR*^C^ 3 h** | **WT vs Δ*tilR*^D^ 6 h** | **Product^E^** |
| --- | --- | --- | --- | --- |
| *mfsX* | RS24900 | -46.80 | -2.29 | Multidrug efflux MFS transporter |
| *uvrX* | RS24895 | -30.24 | -2.47 | Excinuclease ABC subunit UvrA |
| *hmoX* | RS24885 | -4.72 | -1.37 | 4-hydroxyphenylacetate 3-hydroxylase family protein |
| *adsX* | RS24880 | -6.02 | -1.53 | Anthranilate synthase family protein |
| *icmX* | RS24875 | -5.41 | -1.44 | Isochorismatase family protein |
| *dhbX* | RS24870 | -19.32 | -1.50 | SDR family NAD(P)-dependent oxidoreductase |
| *aroX* | RS24865 | -183.92 | -1.45 | 3-deoxy-7-phosphoheptulonate synthase |
| *npsA* | RS24860 | -43.62 | -1.38 | Amino acid adenylation domain-containing protein |
| *thdA* | RS24855 | -19.49 | -1.52 | Acyl carrier protein |
| *npsB* | RS24850 | -4.25 | -1.38 | Non-ribosomal peptide synthetase |
| *npsC* | RS24845 | -4.68 | -1.29 | Nuclear transport factor 2 family protein,  polyketide antibiotic cyclase^F^ |

**^A^**^,^**^B^** ^and^ **^E^** Gene names, locus tags (abbreviated from NDX50_RSXXX) and gene products as annotated for *K. oxytoca* AHC-6 (GenBank accession no. NZ_CP098757.1).

**^C^** ^and^ **^D^** Fold changes in transcript levels of *K. oxytoca* AHC-6 WT compared to Δt*ilR* at 3 and 6 h in LBG. Fold changes were significant (*q* < 0.05) at 3 h whereas they were not significant at 6 h. **^F^** Product prediction by InterProScan.

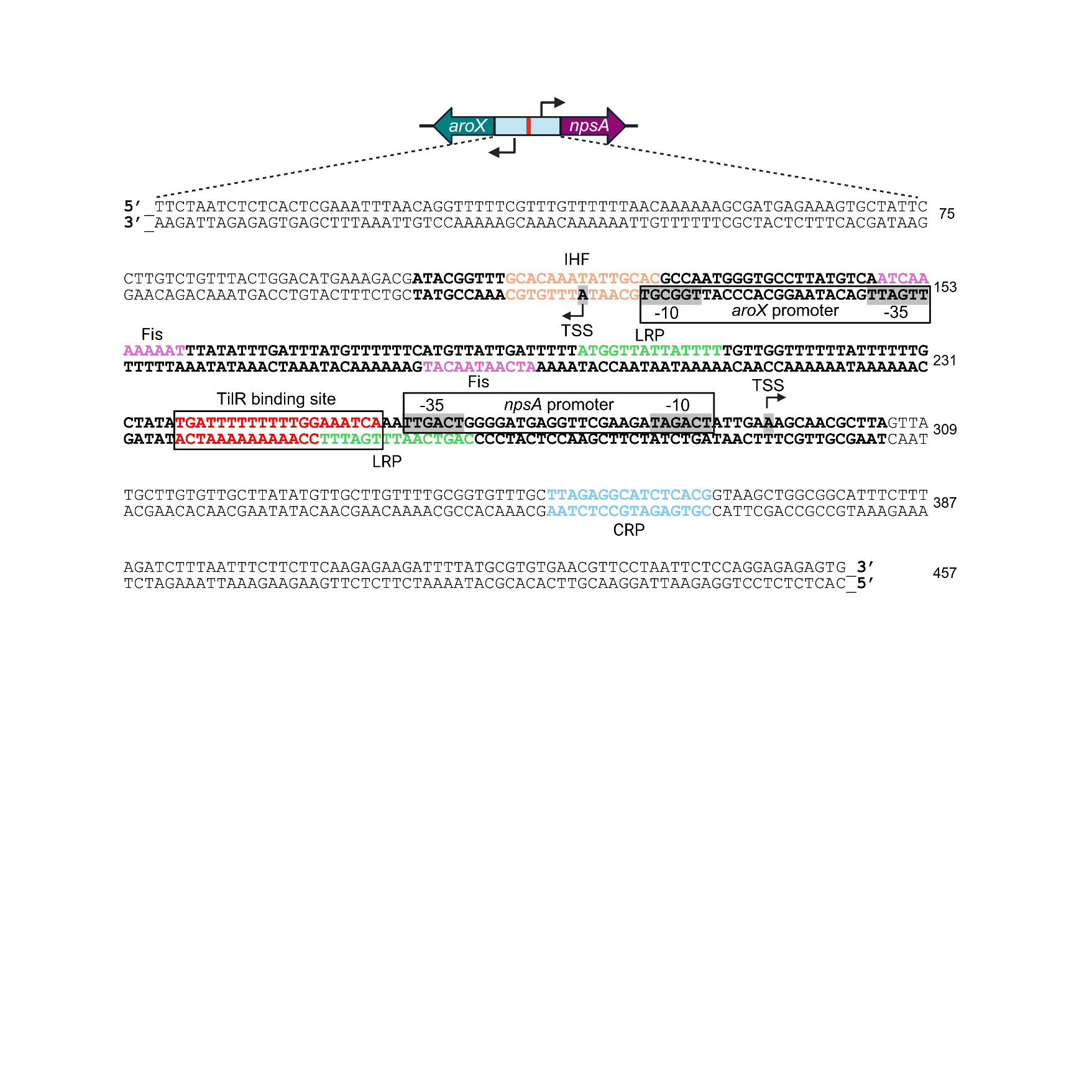
**Figure S4. The *aroX*-NRPS intergenic region.** The sequence is shown in 5’-3’ and 3’-5’ orientations, with nucleotide positions indicated on the right. Binding sites for the transcriptional activators Fis, IHF, LRP, CRP (3-5) and repressor TilR, as well as the candidate promoters (-10, -35 region and TSSs) for *aroX* and *npsA,* are labeled.

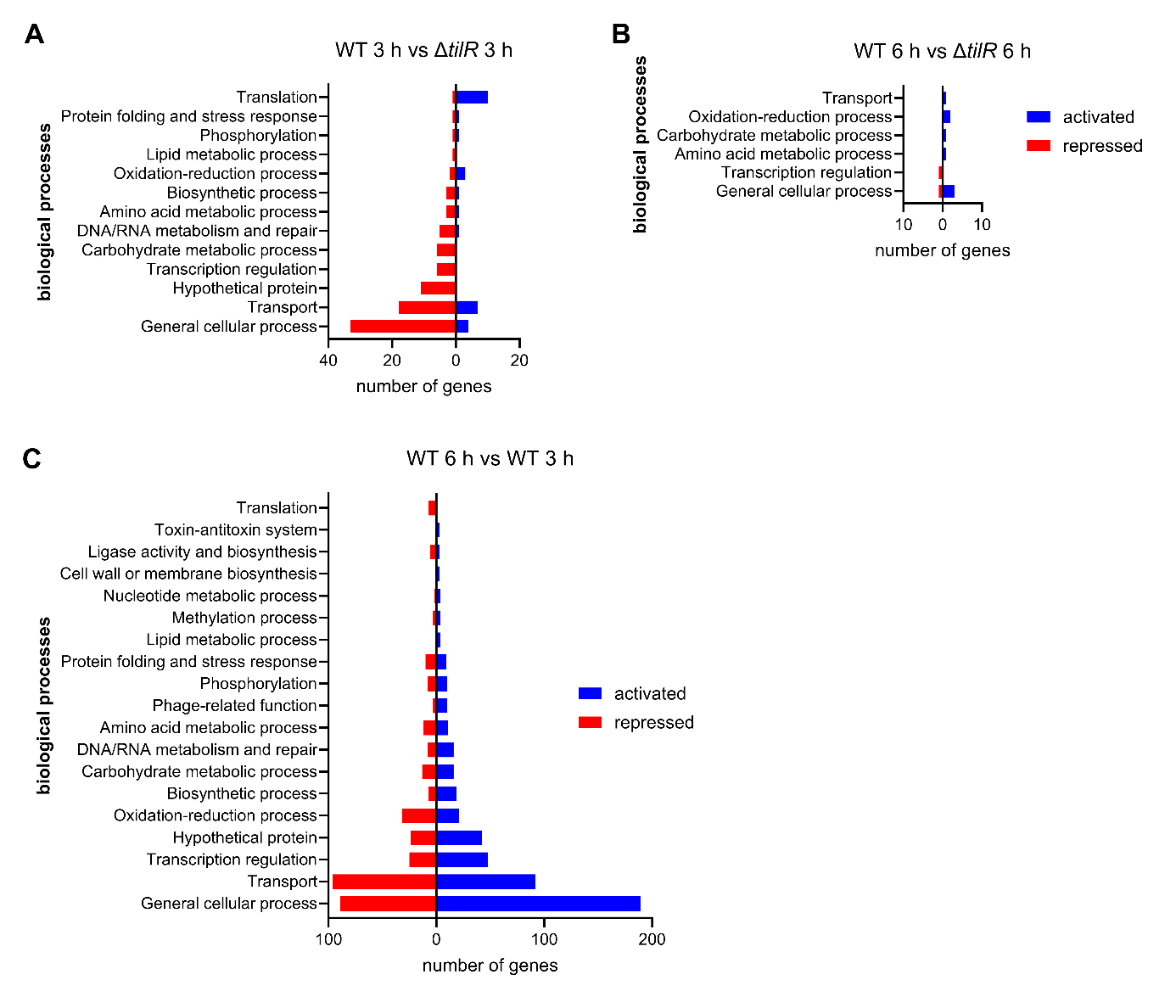
**Figure S5**. **Functional categorization of** **differentially expressed genes in *K. oxytoca* AHC-6.** Differentially expressed genes identified in WT vs. Δ*tilR* at 3 h **(A)** and 6 h **(B)**, and in WT at 6 h vs. 3 h **(C)**, were assigned to biological processes based on Gene Ontology (GO) terms determined using InterProScan. Bars indicate the number of genes expressed at higher (blue) or lower (red) levels in the first condition relative to the second for each comparison.

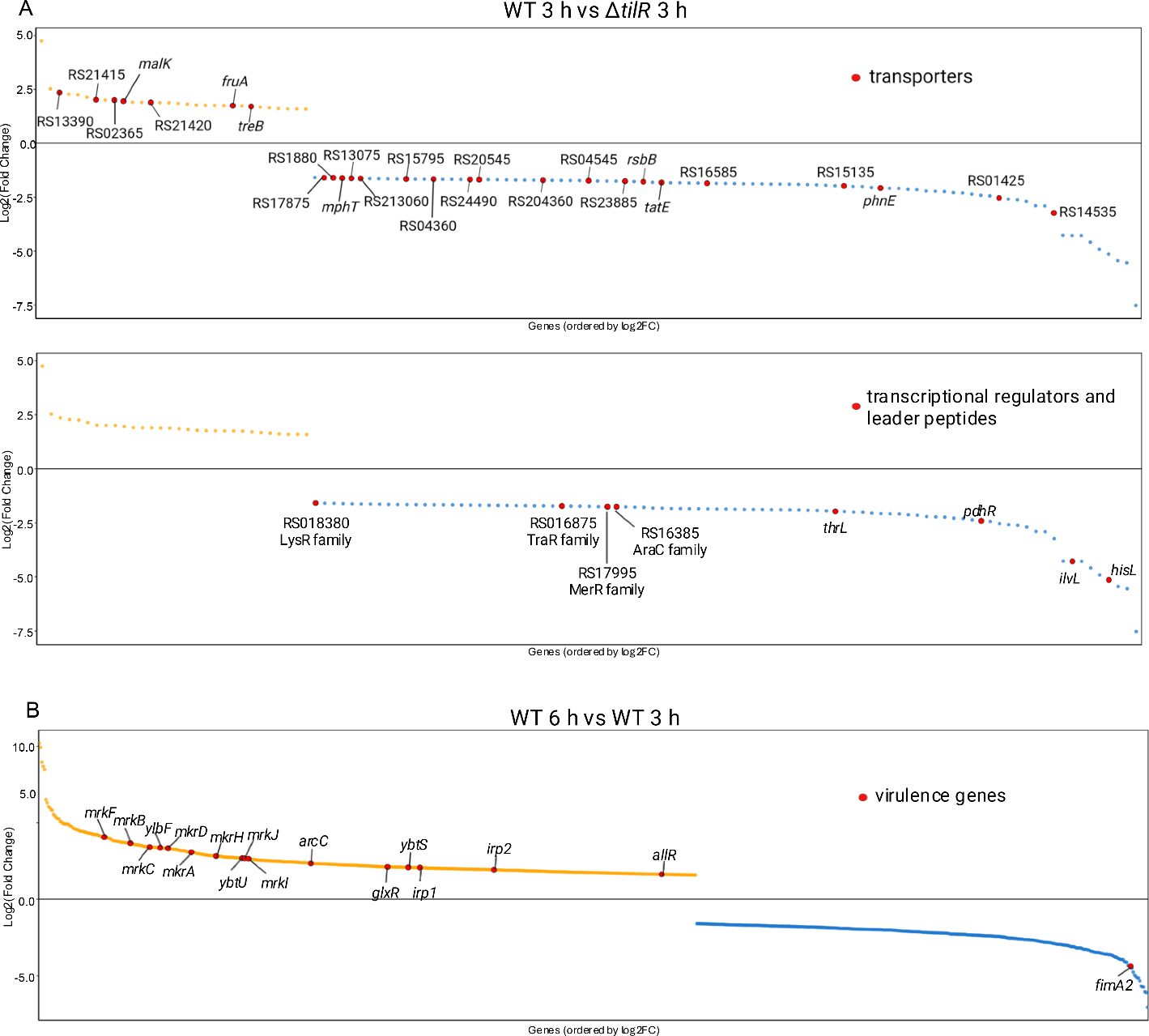
**Figure S6. Notable genes differentially regulated in WT and Δ*tilR* strains*.* (A)** RNA-seq analysis reveals genes associated with transport and transcriptional regulation were among those differentially expressed in WT 3 h vs Δ*tilR* at 3 h. **(B)** Repressed and activated virulence-associated genes in WT 6 vs WT 3 h. The gene products and predicted functions corresponding to the abbreviated locus tags for transporters are provided in the main text or in **Table S4**. Activated genes are highlighted in yellow, repressed in blue, and notable genes are labeled in red.

**Table S5**. Differentially regulated genes in WT vs Δ*tilR* with potential upstream TilR binding sequences.

| **Gene name/**  **product^A^** | **Locus tag^B^** | **Predicted function^C^** | **St.^D^** | **Score** | **p-value** | **Fold change** | **Matched sequence^E^** |
| --- | --- | --- | --- | --- | --- | --- | --- |
| **TGATTTNNNNNNNNMAATYA (6 bp)** | | | | | | | |
| *acpP* | RS18435 | Fatty acid/polyketide synthesis | - | 20.11 | 7.82E-08 | -3.15 | TGATTTACTGAAGCCAATCA |
| *malK* | RS09260 | Transport of maltose/maltodextrin | - | 14.86 | 4.59E-06 | 3.78 | TGATGTCGCCTGCAAAATTA |
| Alginate lyase protein | RS24070 | Digestion of alginate | + | 14.12 | 1.12E-05 | -3.28 | TGATTTTGTTAAAACATTCA |
| Hypothetical protein | RS03880 | - | - | 10.86 | 1.63E-05 | -3.77 | TGAATTGCCGCCTTCAATGA |
| SDR family NAD(P)-dependent oxidoreductase | RS01310 | Short-chain dehydrogenase reductase | - | 10.61 | 1.86E-05 | -3.30 | TGATTTGGATGACCTAAGCA |
| *tatE* | RS15895 | Sec-indep. protein translocase protein TatA/E, part of Tat system | + | 10.17 | 2.76E-05 | -3.53 | TGATTTTCGCGGAGCTTTCA |
| YciN family protein | RS19300 | - | + | 9.88 | 4.62E-05 | -7.44 | TTATTCGGCGCCTGCAATTA |
| Hypothetical protein | RS14960 | - | + | 9.66 | 5.88E-05 | -3.57 | GGATTATGCTCAGACAATCA |
| Hypothetical protein | RS24370 | - | + | 9.66 | 5.88E-05 | -3.18 | TGCTTTAGCTATCGCCATCA |
| DNA-directed RNA polymerase subunit alpha | RS05335 |  | + | 9.64 | 7.37E-05 | 3.13 | TGATCTGTCTGCGGACATTA |
| *rplF* | RS05385 | 50S ribosomal protein L6 | + | 9.64 | 7.37E-05 | 3.52 | TGGTCTTGGTGGCGAAATTA |
| **TGATTTYTTTTTKGMAATYA (20 bp)** | | | | | | | |
| Nitrous oxide-stimulated promoter family protein | RS01795 | - | - | 11.64 | 4.37E-06 | 4.00 | TGATTATTTTGTTGCAAACG |
| Fe-S-containing protein | RS23035 | Membrane iron-sulfur containing protein FtrD-like | + | 7.09 | 2.13E-05 | 3.06 | ATATTACTTTTTTGTAAGCA |
| Nucleobase:cation symporter-2 family protein | RS17875 | Xanthine/uracil permease | - | 6.81 | 2.71E-05 | -3.03 | TTATTCATTTTTTTCAAACA |
| Fimbria/pilus periplasmic chaperone | RS24960 | Pili assembly chaperone | - | 6.55 | 3.18E-05 | 3.67 | CTCTTTCTTTTTCAAAATTA |
| *wbaP* | RS25320 | Undecaprenyl-phosphate galactose phosphotransferase | - | 5.87 | 5.42E-05 | 3.34 | ATGTTTCTAATTTGAAATTA |
| *pdhR* | RS12180 | Pyruvate dehydrogenase complex transcriptional repressor | + | 5.59 | 5.79E-05 | -5.32 | TAATTTTTCGTTTCCATTTA |
| CsbD family protein | RS09305 | - | - | 3.79 | 5.88E-05 | -3.57 | AGATTTATTATTCGGAATCT |
| Hypothetical protein | RS14685 | - | - | 2.56 | 8.16E-05 | -3.18 | TGATTTATTTTTGTGCTTTG |
| Hypothetical protein | RS01900 | - | - | 2.28 | 9.08E-05 | -3.86 | GTATTTTTTATTAACAATAA |

Motifs TGATTTNNNNNNNNMAATYA and TGATTTYTTTTTKGMAATYA were used to scan 500 bp upstream of differentially regulated genes in WT vs Δ*tilR* at 3 and 6 h using the FIMO online tool.

^A^ Gene names or gene products as annotated in *K. oxytoca* AHC-6 (GenBank accession no. CP098757.1) are shown.

^B^ The locus tags are abbreviated from NDX50_RSXXXX.

^C^ Functions of gene products were predicted with InterProScan.

^D^ Indicates if the motif was matched to the forward (+) or the reverse strand (-).

^E^ The actual matched 20 bp DNA sequence located upstream of indicated gene.

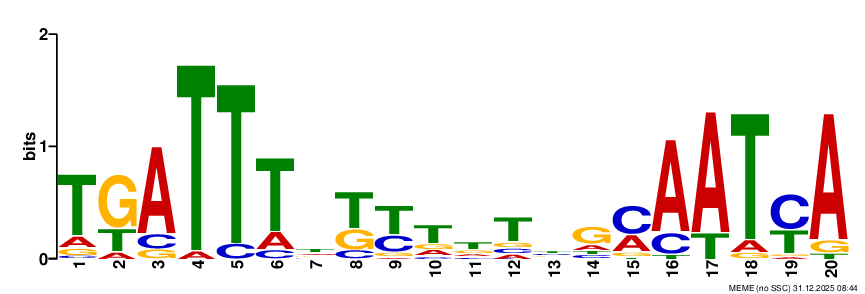

**Figure S7**. **Predicted consensus TilR motif**. MEME was used to create a TilR motif from the 22 binding sites identified in differentially regulated genes (**Table S5**).

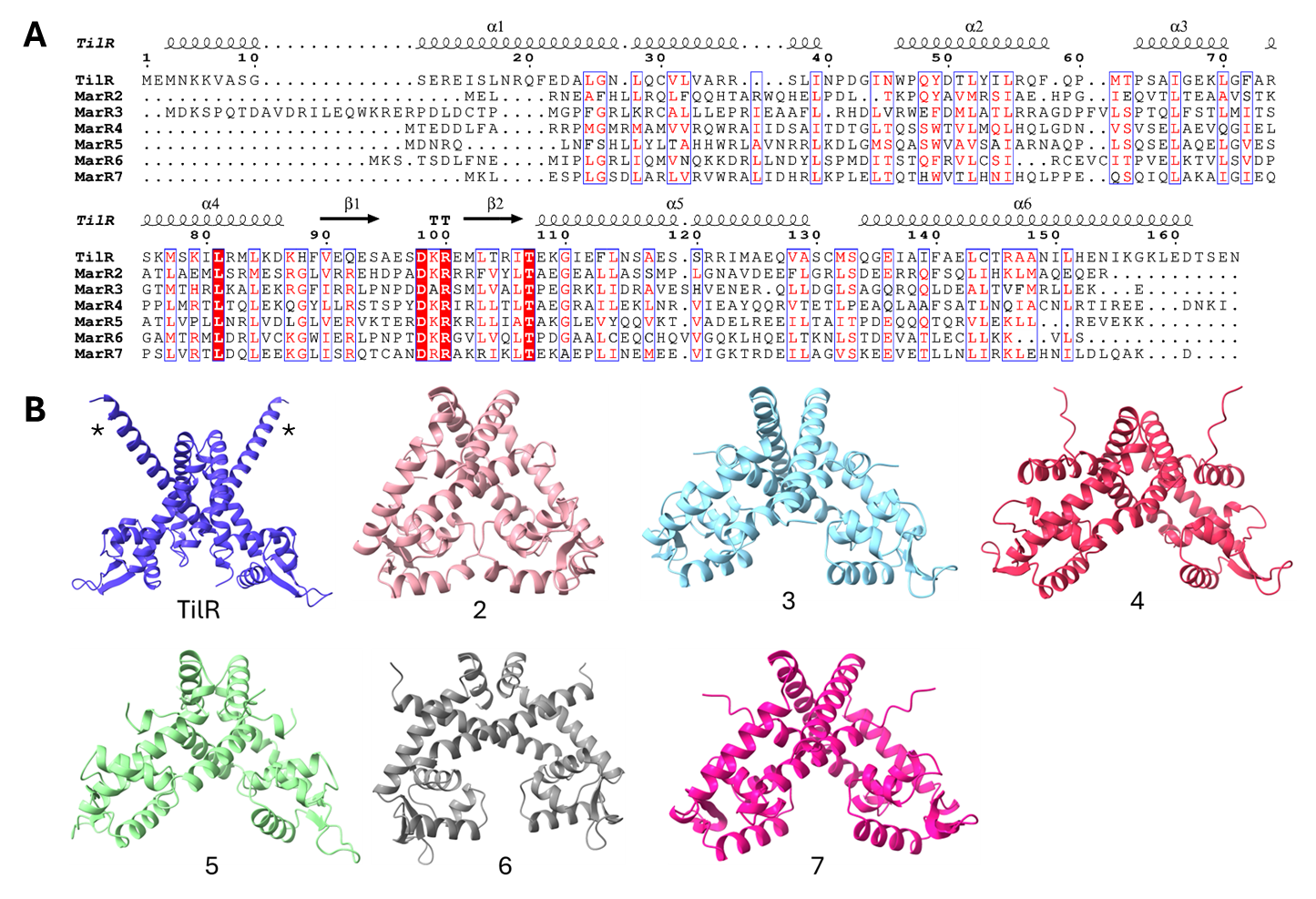
**Figure S8. Sequence and structural comparisons of the seven MarR paralogs in *K. oxytoca* AHC-6 demonstrate TilR is distinct.** (**A**) Protein sequence alignment of the seven MarR-like transcription factors in *K. oxytoca* AHC-6 (performed using Clustal Omega and visualized using ESPript 3.0). Similar residues are marked in red letters outlined with blue boxes, with highly conserved amino acids highlighted in red. Conserved residues are located primarily within the wHTH DNA-binding motif (D98, K99 and R100 in the loop, L81 in α4, and T107 in β2). Secondary structure predictions for TilR and other *Ko* MarR paralogs are indicated above the sequence alignments. The locus tags for *marR*2-7 are, respectively, NDX50_RS01980, NDX50_RS02865, NDX50_RS10935, NDX50_RS20140, NDX50_RS21835 and NDX50_RS22635. (**B**) AF3-predicted structures of *K. oxytoca* AHC-6 TilR (α1 and α1’ are marked with asterisks) and MarR2-7 dimers.

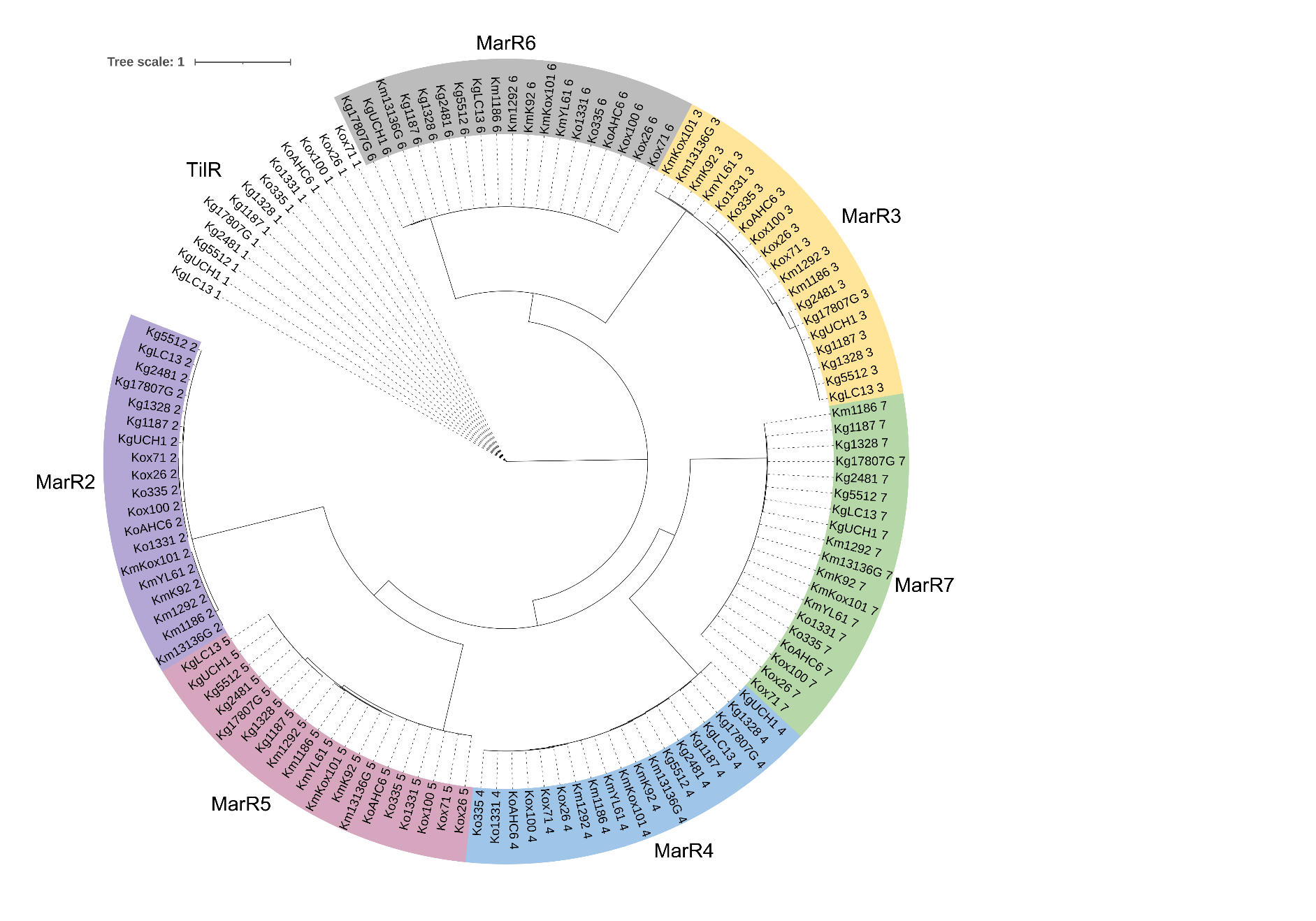

**Figure S9. Prevalence of TilR and its paralogs MarR2-7 in clinical KoSC isolates.** MarR2-7 paralogs are present in *K. oxytoca* (Ko)*, K. grimontii* (Kg)*,* and *K. michiganensis* (Km). TilR is present in Ko and Kg but absent in Km. The labels for each isolate contain the strain name and NCBI accession no. are as follows. *K. grimontii* strains UCH1 (GCA_038185545.1), LC-1303-2020 (CP091752), 5512.56 (CP067391), 2481359 (CP067380), 17807G (SAMN15690275)*; *K. oxytoca* strains Kox71 (CP089392), Kox26 (CP089399), Kox100 (CP089411), AHC-6 (CP098757.1), FDAARGOS_335 (CP027426), FDAARGOS_1331 (CP070144); *K. michiganensis* strains 13136G (GCA_038185415.1), 4928STDY7071186 (NZ_LR607342.1), 4928STDY7071292 (NZ_LR607327.1), YL6-1 (CP084415), K92 (CP089315), Kox101 (CP089407). Multiple sequence alignment was performed with MAFFT, following phylogenetic analysis with PhyML 3.0 and tree visualization using Interactive Tree of Life (iTOL). *SAMN15690275 is annotated as *K. oxytoca* in NCBI but was assigned to *K. grimontii* in the present study based on phylogenetic analysis.

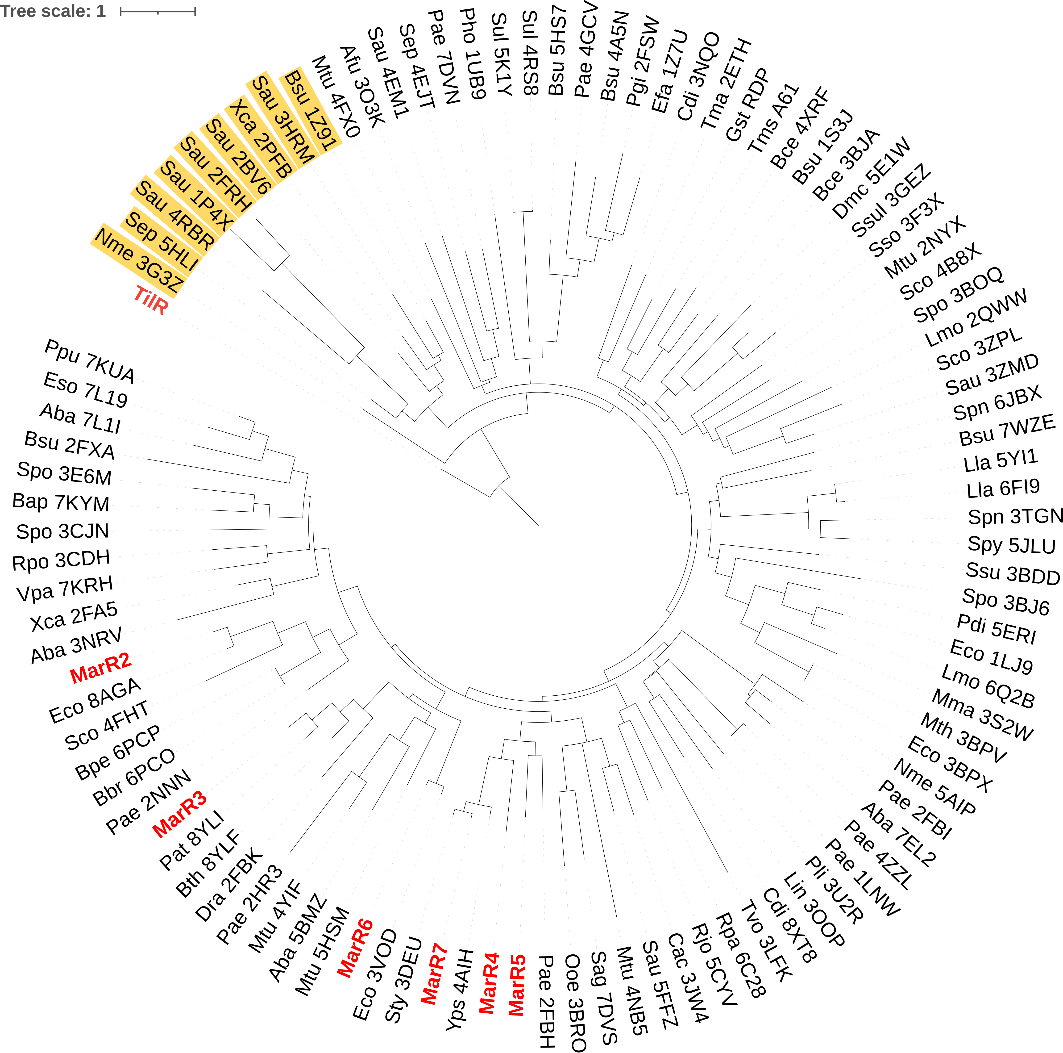

**Figure S10.** **Phylogenetic comparison of TilR and MarR2-7 of *K. oxytoca* AHC-6 with structurally solved MarR-like proteins reinforces the distinctness of TilR**. The Protein Data Bank (RCSB PDB) was searched for MarR-like proteins, yielding 95 solved MarR structures that were used for comparison with MarR paralogs in KoSc. Sequence alignment and analysis were performed using MAFFT and PhyML 3.0, and the resulting phylogenetic tree was plotted with iTOL. *K. oxytoca* AHC-6 TilR and MarR2-7 are indicated in red, while proteins clustering with TilR are highlighted in yellow. The labels for each MarR (95) consist of a three-letter species abbreviation – for example, 'Eco' for *Escherichia coli* – followed by the corresponding PDB code.

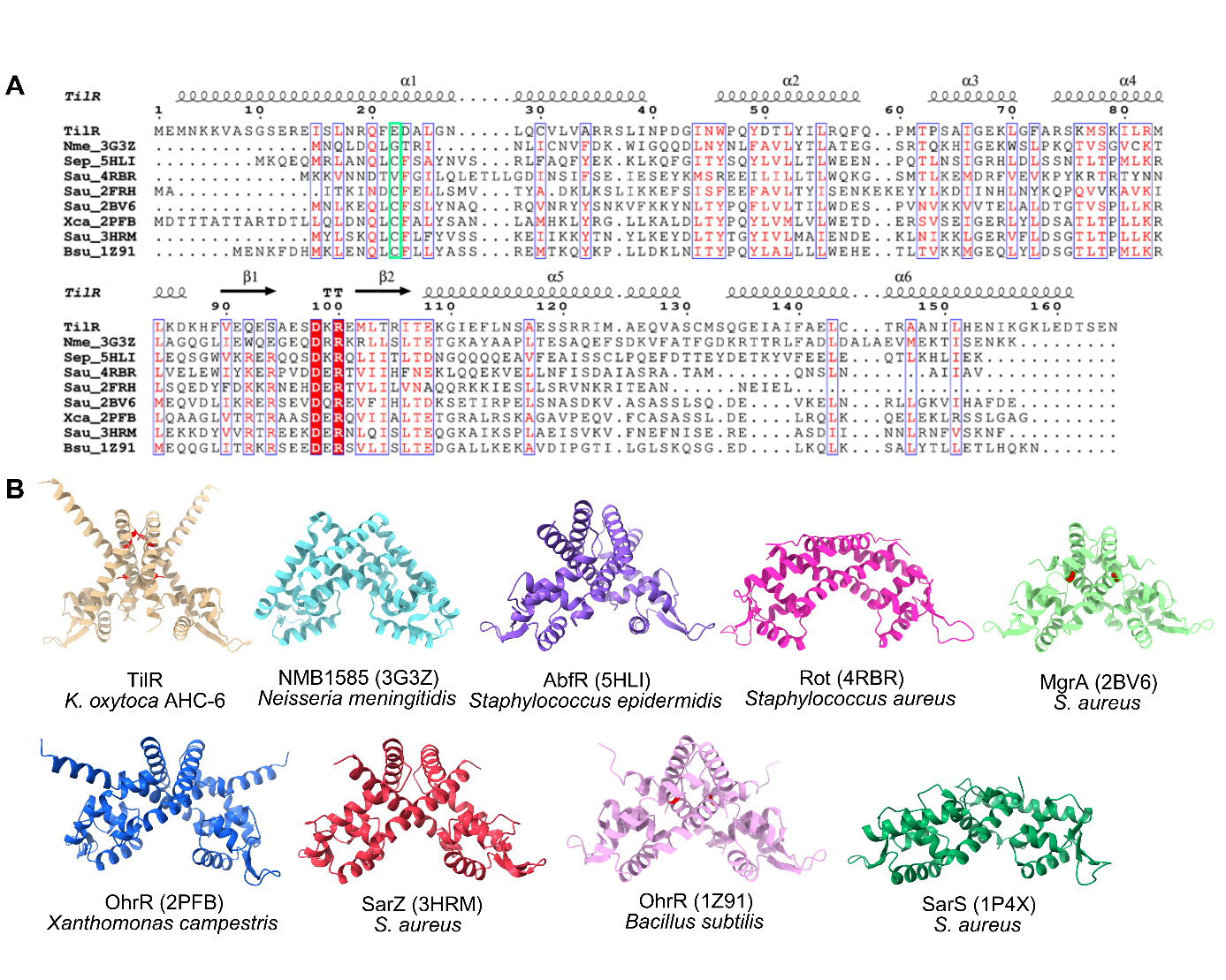
**Figure S11**. **TilR exhibits poor sequence similarity to other homologs and is predicted to possess a distinct MarR-like fold**. (**A**) Multiple sequence alignment of TilR with eight MarR-like transcriptional factors phylogenetically close to TilR identified in **Figure S10**. SarS from *S. aureus* (1P4X) was excluded due to low sequence similarity. Protein sequences of TilR and MarRs with PDB entries 4RBR, 2FRH, 3G3Z, 5HLI, 2BV6, 2PFB, 3HRM and 1Z91 were aligned with Clustal Omega and visualized using ESPript 3.0. Similar amino acids are shown as red letters outlined in blue, with highly conserved residues highlighted in red. Conserved cysteines (C) are marked with a green box. The secondary structure predictions for TilR by AlphaFold 3 are displayed above the sequence alignments. (**B**) Solved structures of the eight MarR dimers. Names of the species and MarR regulator as well as four-letter PDB codes are indicated in parenthesis. Cysteines in TilR, MgrA and OhrR (*B. subtilis*) are displayed in red.

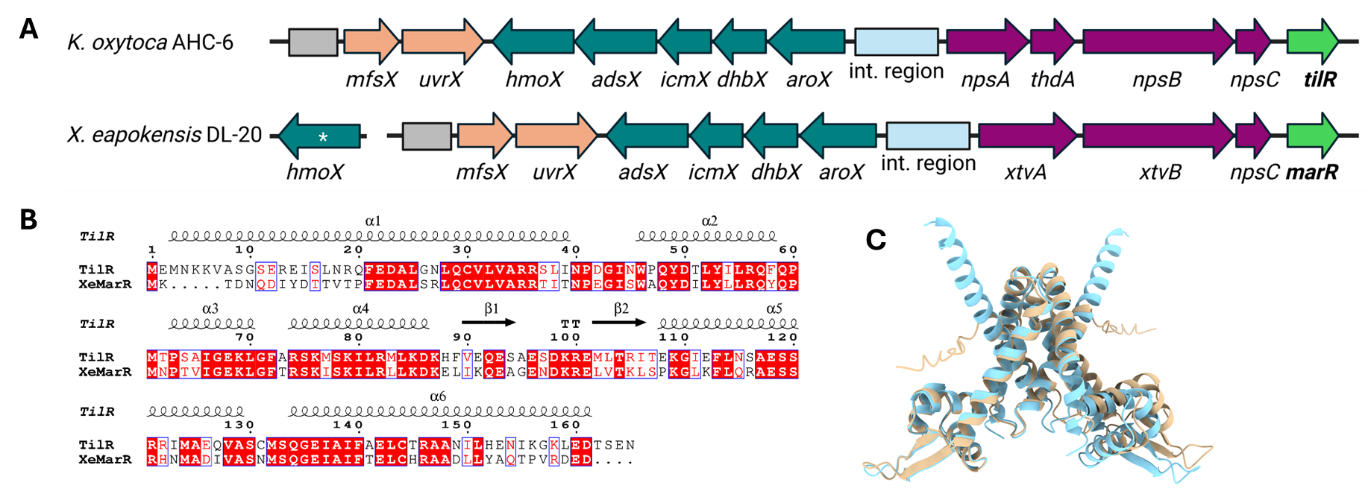
**Figure S12. The *til* gene cluster *in Xenorhabdus eapokensis* encodes a TilR homolog.** (**A**) Comparison of the *til* gene cluster and *mfsX*-*uvrX* operon in *X. eapokensis* DL20 (NZ_MKGQ00000000.1) and *K. oxytoca* AHC-6 demonstrates that all genes are conserved with *xtvAB* equivalent to *npsA*-*thdA*-*npsB*; *hmoX* (asterisk) is encoded elsewhere in the *X. eapokensis* genome. The grey rectangle indicates the upstream putative regulatory region of *mfsX*. (**B**) Protein sequences of TilR and its *X. eapokensis* homolog (XeMarR) aligned with Clustal Omega (65% sequence identity) and visualized using ESPript 3.0 display numerous conserved residues (red boxes). (**C**) Overlay of the AF3-predicted structures for TilR (blue) and XeMarR (beige) displays strong conformational similarity.

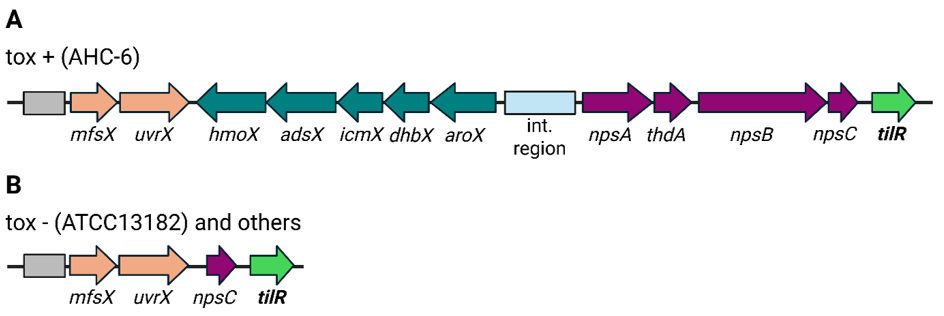

**Figure S13. The *til* gene cluster and *mfsx*-*uvrX* operon in toxin-positive (A) and -negative (B) clinical *K. oxytoca* isolates.** *mfsX*, *uvrX*, *npsC* and *tilR* from *K. oxytoca* AHC-6 (toxin-positive, tox+) are also present in toxin- negative strains (tox-, >99% sequence identity). *K. oxytoca* ATCC 13182 (GenBank accession no. CIP103434, identical to strain NCTC13727), is representative for 14 other clinical isolates including *K. oxytoca* strains KO190245 (CP144730.1), 86 (CP138757.1), FDAARGOS_1333 (CP070020.1), 45 (CP138767), KONSF-1 (CP090245.1), RHBSTW-00373 (CP056567.1), 2024CK-01388 (CP174031.1), 2024CK-01368 (CP174059.1), 2024CK-01266 (CP174080.1), 2024CK-01367 (CP174057.1), A28402 (CP182044.1), A27960 (CP182035.1), 325 (OW849470.1) and FDAARGOS_500 (CP033844.1). The grey rectangle indicates the upstream putative regulatory region of *mfsX*.

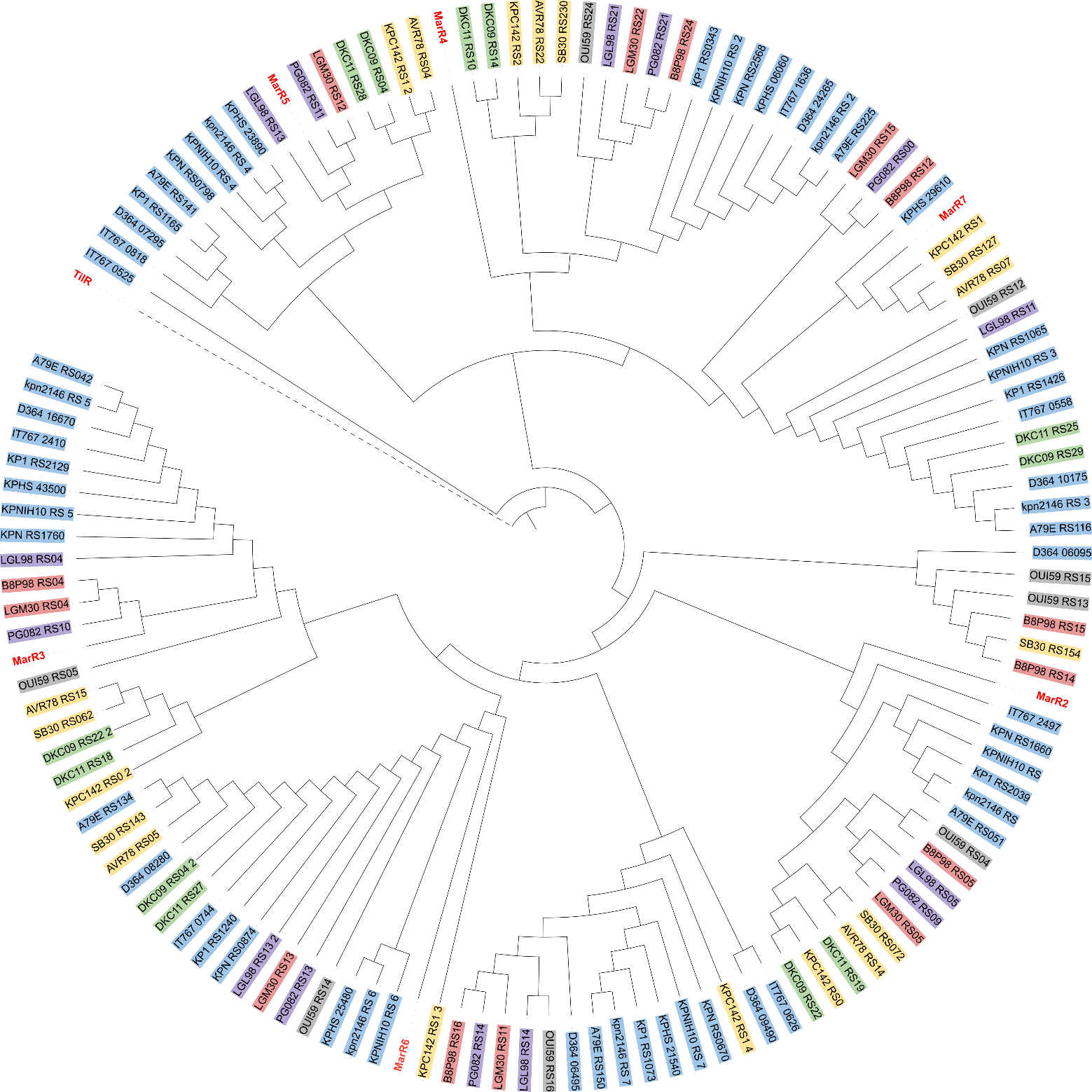
**Figure S14. Conservation of MarR2–7, but not TilR, across representative members of the Klebsiella pneumoniae species complex (KpSC).** MarR2–7 orthologs were identified in 18 representative human isolates of KpSC, including *K. pneumoniae* (Kp1, blue) strains CG43 (locus tags beginning with D364), NTUH-K2044 (KP1), HS11286 (KPHS), ATCC 43816 (IT767), ATCC 700721 (KPN), KPNIH10, 1084 (A79E), and ATCC BAA-2146 (kpn2146); *K. quasipneumoniae* subsp. *quasipneumoniae* (Kp2, green) strains CAV2013 (DKC09) and CAV1947 (DKC11); *K*. *quasipneumoniae* subsp. *similipneumoniae* (Kp4, yellow) strains SB30, ATCC 700603 (AVR78), and KPC142; *K. variicola* subsp. *tropica* (Kp5, gray) strain SB5531 (OUI59); *K. quasivariicola* (Kp6, salmon pink) strains KPN1705 (B8P98) and 08A119 (LGM30); and *K. africana* (Kp7, lavender) strains 200023 (LGL98) and AF01-73J (PG082). Protein sequences were aligned using MAFFT, and phylogenetic analysis was performed using PhyML 3.0, with tree visualization in Interactive Tree of Life (iTOL). Colored label backgrounds distinguish the KpSC phylogroups, while the *K. oxytoca* AHC-6 reference proteins are labeled in red. Orthologs of MarR2–7 were identified across the sampled KpSC strains, whereas no TilR ortholog was detected. The Kp3 phylogroup (*K. variicola* subsp. *variicola*) was excluded because sufficiently reliable genome sequences from human isolates were not identified for inclusion in the analysis.

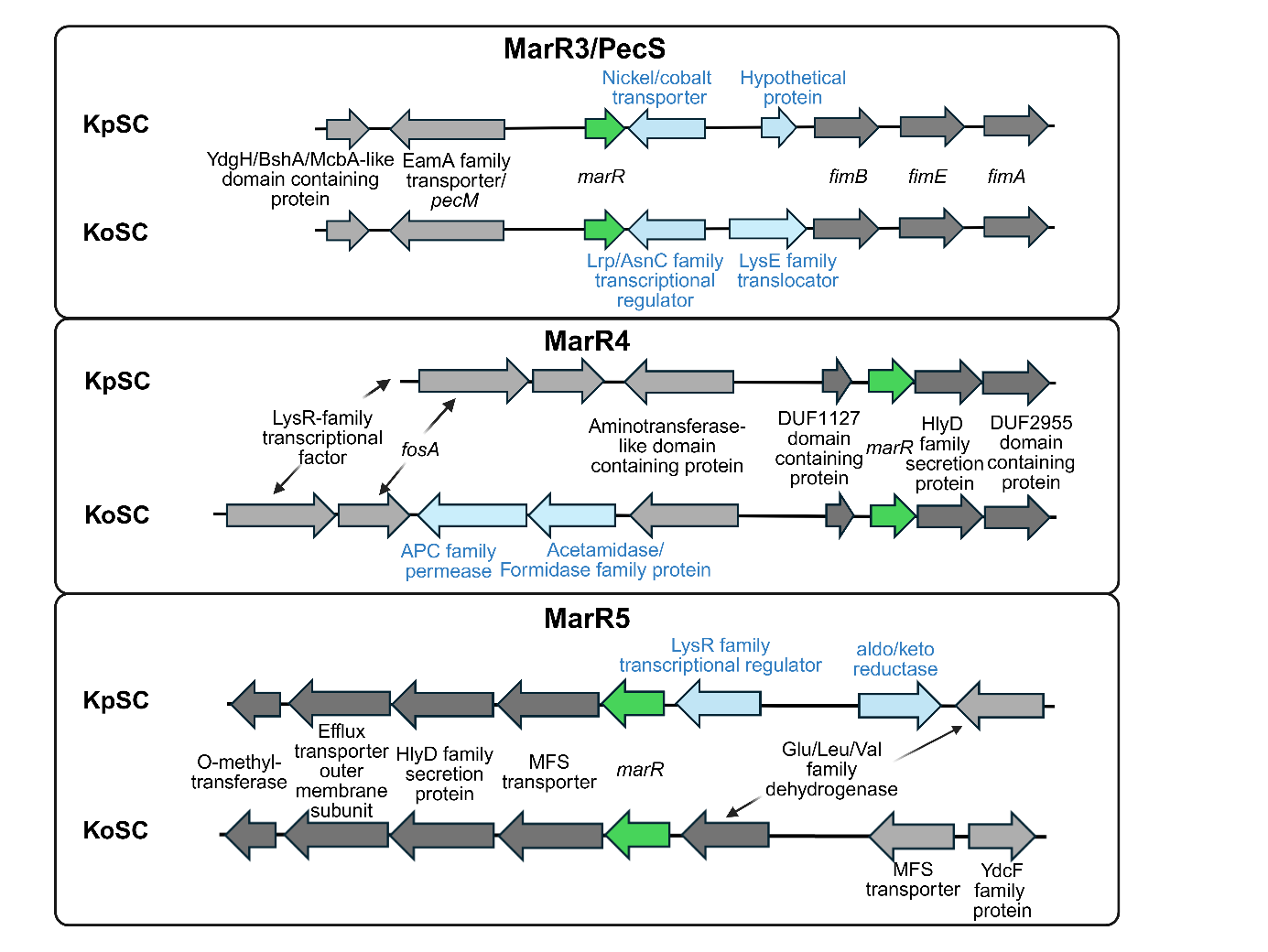

**Figure S15. Comparison of MarR3-5 genetic loci in KpSC and KoSC.** Gene neighborhoods of MarR homologs in *K. pneumoniae* strains CG43 (locus tags starting with D364), NTUH-K2044 (KP1) and ATCC 43816 (IT67) are shown. The flanking genes of MarR3/PecS homologs (KP1_RS21295, D36_16670, IT767_24100), MarR4 (KP1_RS0343, D36_424265, IT767_16365) and MarR5 (KP1_RS11655, IT767_08180, D364_07295) are displayed. Genes and gene products that differ between the *K. oxytoca* AHC-6 and KpSC MarR loci are highlighted in blue; labels for identical genes are positioned between the two loci with arrows for easier orientation. The PecS locus in KpSC contains two genes replaced by a nickel/cobalt transporter and a hypothetical protein. The MarR4 locus additionally encodes an APC family permease and an acetamidase/formamidase family protein, while the MarR5 locus contains a LysR-family regulator and an aldo/keto reductase.
